# Body size, dental pathology and maternal genetic diversity of ancient horses in the eastern Baltic Sea region and western Russia

**DOI:** 10.64898/2026.03.17.712305

**Authors:** Johanna Honka, Daniela Salazar, Arthur O. Askeyev, Igor V. Askeyev, Oleg V. Askeyev, Jouni Aspi, Gulshat Sh. Asylgaraeva, Markku Niskanen, Kristiina Mannermaa, Suvi Olli, Noora Piipponen, Giedrė Piličiauskienė, Dilyara N. Shaymuratova, Renat R. Valiev, Laura Kvist

## Abstract

The early evolutionary history of modern domestic horses (*Equus caballus/E. ferus caballus*), known as the DOM2 lineage, is well documented due to numerous archaeological and ancient DNA (aDNA) studies. Although many uncertainties remain in the domestication timeline, current evidence suggests that the domestication of modern horses began in the Pontic-Caspian steppe at least ∼2700 BCE (before common era), or even earlier. However, it is not known how long remnant wild horse populations survived or when domestic horses were introduced into Northern Europe. In this study, we review the current knowledge of horse domestication, focusing on Northern Europe. We analysed prehistoric horses from western Russia to assess the body sizes of wild horses from the Ivanovskaya site (5900–3800 BCE) in the Pontic-Caspian steppe, and the body weight of one Lithuanian wild horse (4000-3800 BCE). Additionally, we analysed body sizes of Late Bronze Age-Early Roman Age horses (1100 BCE-300 CE; common era) and re-analysed body sizes and estimated rider weights of historic domestic horses from Lithuania (100-1400 CE). We searched for pathological changes and signs of bit wear indicative of bridling. Furthermore, we investigated maternal genetic diversity by sequencing ancient mitochondrial DNA. We found that wild horses from Ivanovskaya were intermediate in body size between earlier and more recent horses of the Eurasian Steppe, and that the Lithuanian wild horse weighed only ∼270 kg and Late Bronze Age-Early Roman Age horses 200-300 kg. Lithuanian domestic horses were pony-sized (< 130 cm on average). Bit wear was confirmed on one tooth, the oldest domestic horse in Lithuania (799-570 cal BCE). Another tooth showed signs of the Equine Odontoclastic Tooth Resorption and Hypercementosis (EOTRH) condition. Mitochondrial DNA was successfully amplified from one Ivanovskaya wild horse along with 25 other ancient samples, including Lithuania’s oldest domestic horse. mtDNA diversity was high, revealing several maternal lineages.

## Introduction

Horses first appeared in the fossil record ∼50 million years ago, and over millions of years these early horses evolved from dog-sized mammals into the single-toed genus *Equus*, and eventually into the horses we know today. Wild horses (*E. ferus or E. f. ferus*; Fig. 1) were once abundant in the late Pleistocene mammoth steppe, but as their prime habitat began to disappear during the Pleistocene-Holocene transition (∼9600 cal BCE; before common era), their populations declined. Based on archaeological evidence, wild horse population sizes fluctuated during the Holocene (Petrenko, 2007; Sommer et al., 2011; Leonardi et al., 2018), with free-living horses in Central Europe and the Baltics observed until the end of the 16^th^ century CE (common era) (Lappo, 1938; van Vuure, 2014). However, it is unknown when exactly wild horses went extinct. European wild horses, known as Tarpans, are claimed to have persisted until the 19^th^ century; however, their late existence is not supported by tangible scientific evidence, historical sources or DNA (Lovász et al., 2021, Librado et al., 2021; 2024). At the time of horse domestication, two additional horse lineages existed: the Lena horse (*E. f. lenensis*) in Siberia and an Iberian horse population (IBE). Both are now extinct and did not significantly contribute to the ancestry of the modern domestic horse lineage (Fages et al., 2019; Lira Garrido et al., 2025).

**Fig. 1.**
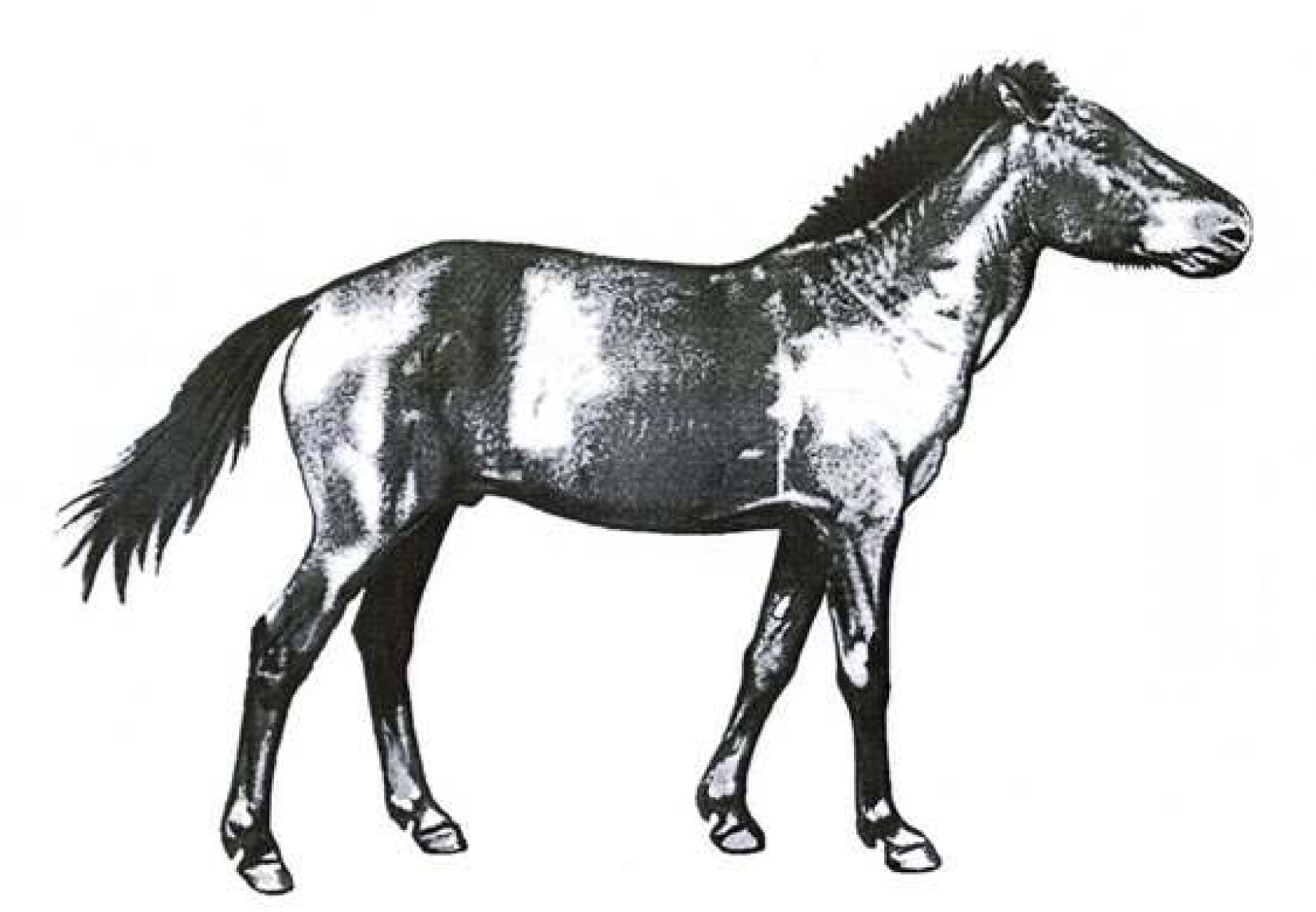
Reconstruction of a late Palaeolithic/late Pleistocene wild horse stallion (*Equus ferus* or *E. ferus ferus*), based on four rock paintings from the Shulgan-Tash (Kapova) cave in the Southern Urals, Republic of Bashkortostan, Russia (Ruiz-Redondo et al., 2020). Morphometric data and other morphological features of late Pleistocene Eastern European and Southern Ural wild horses were also used in creating the reconstruction (Kuzmina, 1997; Plasteeva, 2013). Illustration by Igor Askeyev.

Today, only the domesticated form of the horse persists. Domestication refers to a sustained, multigenerational mutualistic relationship between humans and animals, in which humans control the reproduction and care of the animal (Zeder, 2012). The origins of the modern domestic horse (known as the DOM2 lineage of *Equus caballus/E. f. caballus*) have been traced to the lower Don-Volga region of the Pontic-Caspian steppe (Fig. 2) through ancient DNA (aDNA) (Librado et al., 2021), reviewed in Kanne (2025). The genomic data indicated a domestication bottleneck starting ∼2700 BCE, followed by close kin mating and shortened generation times, suggesting reproductive control by humans around ∼2200 BCE (Librado et al., 2024). However, direct archaeological evidence is still lacking. A temporal gap of 500–700 years exists in the genomic studies of Librado et al. (2021; 2024) between the earliest DOM2 horses and their ancestors, and wild horses from the forest-steppe. In addition, studies from southern Asia are still lacking, leaving the precise region of domestication open for debate. DOM2 horses spread beyond the Western Eurasian steppe by 2200-2000 BCE (Librado et al., 2021), eventually replacing all other horse lineages by 1500-1000 BCE, except those ancestral to the historic Przewalski’s horse (*E. (f.) przewalskii*) and the Tarpan (Librado et al., 2021, 2024). DOM2 horses are suggested to have reached eastern and southeastern Mongolia at least by 1400-1300 BCE (Honeychurch et al. 2025). However, these genomic estimates of the domestication timeline carry many uncertainties regarding underlying model assumptions and the fact that the domestication status of the horses cannot be confirmed. Further archaeological and genomic evidence is needed to confirm the chronology of domestication.

**Fig. 2.**
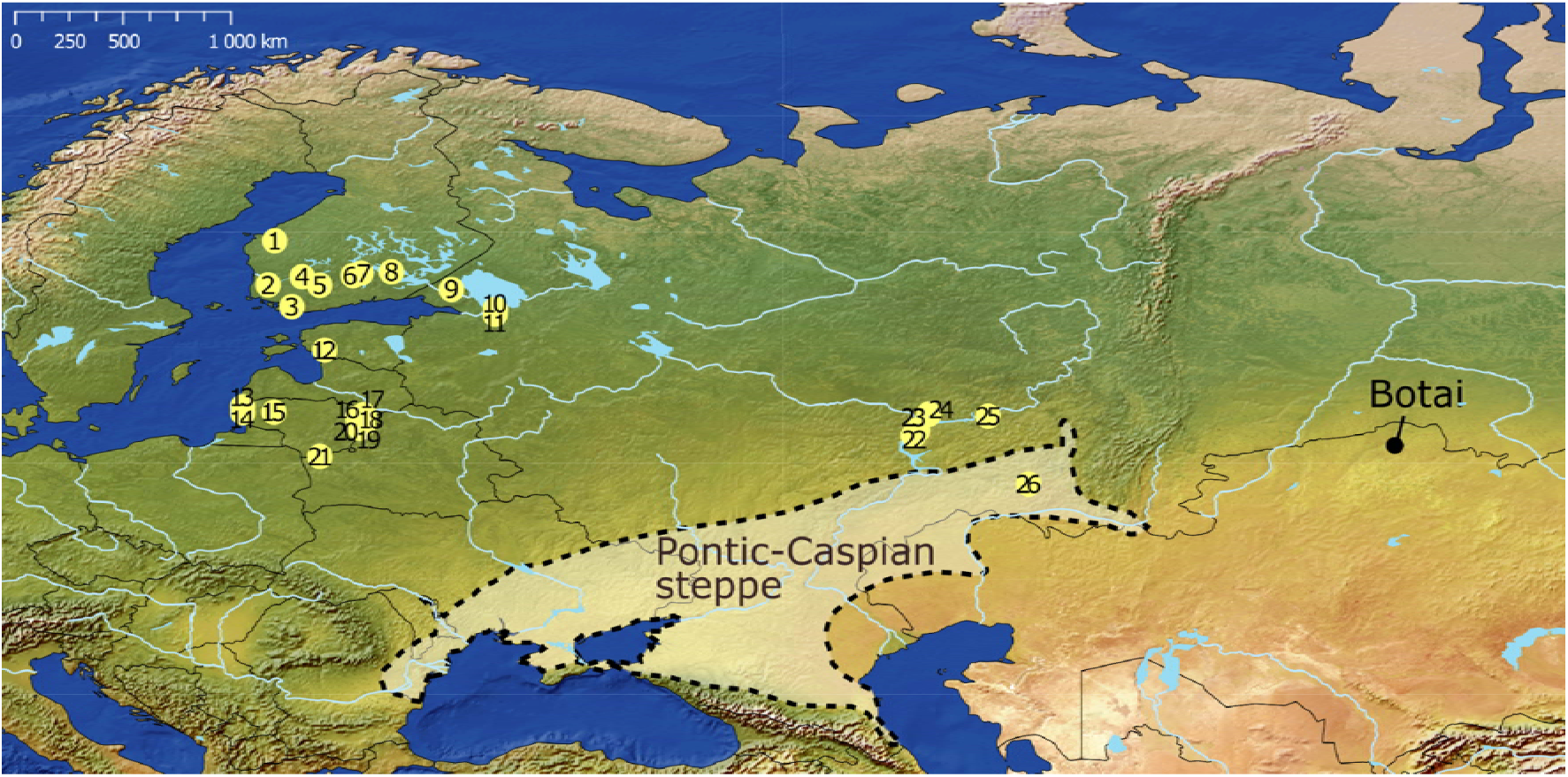
Sampling locations of the archaeological horses in this study are indicated by numbered yellow circles. Sampling sites: 1. Levänluhta water burial, Isokyrö, 2. Eura, Luistari, 3. Knaapila, Salo (Perniö), 4. Tursiannotko, Pirkkala, 5. Linnavuori, Tenhola, Hattula, 6. Ihananiemi, Sysmä, 7. Uusi-Ruskeala, Hartola, 8. Latokallio, Moisio, Mikkeli, 9. Tontinmäki, Hovisaari, Räisälä, 10. Ladoga Canal, 11. Podolije 3, 12. Pärnu, 13. Šventoji 43 (probable wild horse), Šventoji 41B and Šventoji 23, 14. Kukuliškės, 15. Daktariškė 5 (probable wild horse), 16. Garniai 1, 17. Mineikiškės, 18. Antilgė, 19. Kretuonas 1A, 20. Garnys, 21. Kabeliai 7, 22. Second Polyansky burial, 23. Novoslavski II burial, 24. Pestrechinskaya II, 25. Biklyan settlement and 26. Ivanovskaya (wild horse). Sampling sites 1-8 are in Finland, 9-11 and 22-26 in Russia, 12 in Estonia and 13-21 in Lithuania. The origin of modern domestic horses (DOM2) in the Pontic-Caspian steppe is also marked, along with the archaeological site of Botai, which contains the earliest potential evidence of domestic horses (DOM1). As a background raster image, 1:50m Natural Earth II (public domain) was used, portraying the world in an idealized manner as it appeared before the modern era, with minimal anthropogenic influence.

Around the time of the domestication of the ancestors of the DOM2 lineage in the Pontic-Caspian steppe, a genetically distinct population, known as DOM1, is proposed to have been domesticated in the northern Kazakh steppe (Gaunitz et al., 2018; Librado et al., 2021). Today, this lineage persists in Przewalski’s horses. Evidence of horse domestication in this region is primarily associated with the Botai culture (3500-2700 BCE); however, the debate over whether these Botai horses were domesticated or not remains ongoing (e.g. Outram 2023), with the possibility that both wild and domestic horses were present at Botai (e.g. Chechushkov & Kosintsev, 2020; Niskanen, 2023).

Archaeological evidence suggests even an earlier period of horse domestication or taming, dating to the 4^th^ or even 5^th^ millennium BCE. Human skeletal pathologies associated with horseback riding have been identified in pre-Yamnaya and Yamnaya individuals from Romania, Bulgaria and Hungary dating to the 5^th^ - 3^rd^ millennium BCE (Trautmann et al., 2023; the earliest rider from Trautmann et al., 2023 was redated to 4331–4073 cal BCE in Lazaridis et al., 2025; see also Kanne, 2022 for the Hungarian individuals), but riding-related diagnostic traits co-occurred only in a small proportion of the hypothesized riders (Hosek et al., 2024). Horse milk proteins were found in the dental calculus of two early Bronze Age humans (3^rd^ millennium BCE) from the lower Don River estuary in the Pontic-Caspian steppe (the earlier of these individuals was redated to ∼2900 BCE in Lazaridis et al., 2025; milk peptide study in Wilkin et al., 2021). However, Scott et al. (2022) found no evidence of horse dairying before the 9^th^ century BCE in the North Caucasus steppe region, suggesting horse milking had been a very limited activity. Overall, due to a lack of direct evidence and different interpretations of riding-related skeletal changes, the evidence of horse domestication remains inconclusive (Niskanen, 2023).

Farming and animal husbandry started in Fennoscandia (Sweden, Norway, Finland, Russian Karelia and Russian Kola Peninsula) and the Baltic countries (Estonia, Latvia and Lithuania) at the 4^th^ millennium BCE (Vanhanen et al., 2019) and at the 3^rd^ millennium BCE (Oras et al., 2023), respectively. The timing of horse introduction to the Baltic states and Fennoscandia remains unknown. The earliest horse remains in Lithuania were found at the Daktariškės 5 and Šventoji 43 sites (see Fig. 2), dated to 4540-4369 cal BCE and 3958-3798 cal BCE, respectively (Honka et al., 2025), indicating they were likely wild. In Lithuanian Subneolithic (4500-2900 BCE) and Neolithic (2900-1800 BCE) sites, horse remains are scarce, comprising up to 1-2% (Piličiauskas et al., 2023) of the recovered remains. Between 1100-500 cal BCE, the number of horse remains increases considerably, especially at sites in western Lithuania and western Latvia, where they account for 10-40% of all mammal remains (Bliujiene et al., 2020; Piličiauskas et al., 2023; Vasks et al., 2011; 2020). In Estonia, the earliest domestic horse has been dated to 700-500/400 cal BCE (Librado et al., 2021; reviewed in Rannamäe et al., 2024). Similarly, in Finland, the earliest horse bone was radiocarbon dated to 820-546 cal BCE (Bläuer & Kantanen 2013). In Scandinavia (Sweden, Norway and Denmark), the first horse bones were dated to ∼1600 cal BCE (Kveiborg, 2018; Kveiborg et al., 2018). Lithuanian and Estonian horses (dates ranging from 800 BCE to 1800 CE) were found to be small, usually in the range of ponies, with average withers heights ranging from 122-132 cm across different sites (Bliujiene et al., 2017; Piličiauskienė et al. 2022; Rannamäe et al., 2025), with the body mass of Estonian horses averaging 288 kg (Rannamäe et al., 2025). The body sizes of ancestral horses of the Pontic-Caspian steppe are unknown, and the estimations of withers height for Lithuanian horses follow May (1985), which does not take differences in limb segment proportions into account. Additionally, rider weights estimated from skeletal dimensions have not been calculated for the small Lithuanian horses. Ancient DNA and multidisciplinary research are needed to confirm the arrival times of domestic horses in Fennoscandia, the Baltic states and western Russia, as differentiating wild from domestic horses based on the morphology of archaeological samples is challenging.

Further uncertainties remain as to whether the genetic lineages of early horses survive today in landrace horse breeds or whether they were replaced by later waves of domestic horses. Several native northern horse breeds currently exist in the Baltic states and Fennoscandia, such as the Estonian Native Horse, the Žemaitukas horse in Lithuania, the Finnhorse in Finland, the North Swedish horse and the Gotland Russ Pony in Sweden, the Fjord Horse and the Nordland/Lyngen horse in Norway, and the Icelandic horse, which could descend from the early domestic horses brought to these areas. Skeletal measurements of Estonian archaeological horses and modern Estonian Native horses imply that the horses are of a similar type, although their size has increased and their limbs have become more slender (Rannamäe et al. 2025). The territory of present-day Russia also harbours many native breeds, such as the indigenous Tuva, Vyatka and Yakut horses, as well as the Don, Kabardin and Akhal-Teke horses. Many cold-blooded native horse breeds in these countries share genetic similarities, indicating a common history and admixture among these breeds, as well as genetic affinities to eastern breeds and even to Przewalski’s horse (Castaneda et al., 2019; Kvist et al., 2019; Sild et al., 2019; Kvist & Niskanen, 2021).

While the domestication history of horses has been extensively studied through aDNA, the spread of domestic horses to northern areas and the characteristics of the horses are poorly understood. Moreover, historical records of domestic horses in the eastern Baltic Sea region are scarce, primarily from ∼1200-1800 CE; thus, understanding their history and prehistory relies heavily on subfossil, i.e. archaeological finds (Bläuer et al., 2022). Additionally, the body size of the ancient Northern horses, except those from Estonia, or the wild horses from the Pontic-Caspian steppe has either not been studied using skeletal measurements or has been studied only using older, less-accurate methods. Dental changes related to bridling (bit wear) are informative about the use of the horse for riding, while signs of infection indicate horse health. Therefore, we used a multidisciplinary approach to study the early history of horses of the eastern Baltic Sea region and western Russia, spanning from wild horses to domestic horses from later times. Our aims were to 1) evaluate the body sizes of wild horses from a site within the Pontic-Caspian steppe, ancestors to DOM2 horses, as well as temporal changes in body size of domestic horses from Lithuania using skeletal measurements and updated size estimation methods. Additionally, we estimated rider weights and compared them to the sizes of average contemporaneous humans. We also 2) investigated possible pathological changes in our samples to identify how humans utilized the horses, whether the horses suffered any illnesses or infections, and to possibly differentiate wild horses from domestic horses through dental changes associated with bridling. In addition, we 3) studied maternal genetic lineages shared among different archaeological sites and countries by analysing a 572–573 bp (base pair) fragment of the mitochondrial control region from ancient DNA extracted from skeletal samples to assess regional differences.

## Material and methods

### Body size estimations

We assessed body sizes (withers height, croup height and body mass) for 25 wild horses from the Ivanovskaya site (Russia) based on 25 metapodial and phalangeal specimens dated to 5900–3800 BCE and from two phalangeal (Ph1 forelimb and hindlimb) specimens from Lithuanian Late Bronze Age (approx. 1100–500 BCE) horse/s (possibly the same individual) from Mineikiškės. Additionally, we re-analysed 43 probable domestic horse metapodials from several Lithuanian sites divided into three temporal periods: Lithuanian pre-Viking Iron Age (∼100-700 CE; *n* = 8), Lithuanian Viking period (∼800-1100 CE; *n* = 9), and Lithuanian Medieval period (∼1200-1400 CE; *n* = 26), using updated body size estimation equations (Table S1-S4). The Lithuanian horses from pre-Viking Age onwards were previously analysed in Piličiauskienė et al. (2022) using equations from May (1985).

For the Ivanovskaya wild horses, the height estimates were adjusted using reference data for both domestic and Przewalski’s horses, as it is not known whether these wild horses, living in the easternmost regions of the Pontic-Caspian Steppe, were more similar to domestic or Przewalski’s horses in the height – bone length relationship (see Method S1, Table S5). For the Lithuanian horses from pre-Viking Age onwards, we additionally estimated the maximum rider weights (XRWs), which include both rider and riding gear, by applying equations from Niskanen (2023: Supplementary Information B), as described in Methods S1 and Tables S6-S8. Calculations of percent prediction error (%PE) and mean ratios between observed or reconstructed heights and estimated heights are provided in Tables S9-S10. The rider weights were compared with estimated sizes of contemporaneous male Lithuanians (Jankauskas, 2001). One of these sites (Marvelė) includes size estimates of both horses and humans. The estimated withers and croup heights and body masses of the Ivanovskaya wild horses were compared with those of the Eneolithic horses from Deriivka (a Sredny Stog culture site in Ukraine) and the Botai and the Tersek culture sites of Kazakhstan (Niskanen, 2023), by re-estimating their metacarpal dimensions using equations introduced in this study (Methods S1).

Additionally, we estimated only body weights of a Lithuanian wild horse from Šventoji 43 site (aEca15; 3958-3798 cal BCE; Table 1) using the left distal tibia, of a Lithuanian horse from Antilgė site from Phalanx I (XX) (aEca4; 800-500 BCE or 100-200 CE; Table 1) and of another horse from Antilgė site based on distal tibia (Late-Bronze-Early Roman Age, ∼1100 BCE-300 CE), using equations in Methods S1. From these bones, the withers and the croup heights could not be estimated. Body size estimations could not be performed for other bones in Table 1 due to incomplete preservation or the lack of suitable reference data.

**Table 1.**
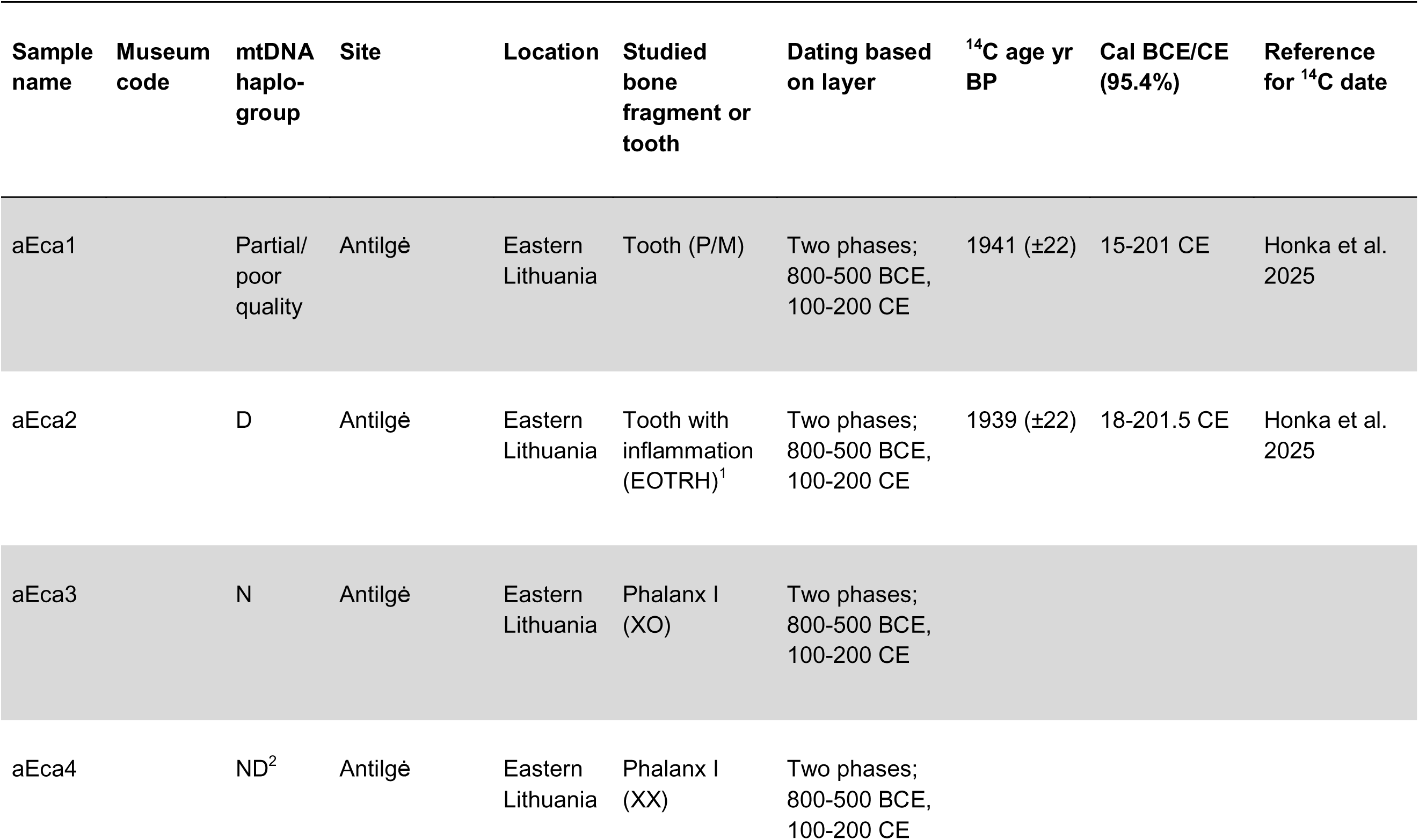
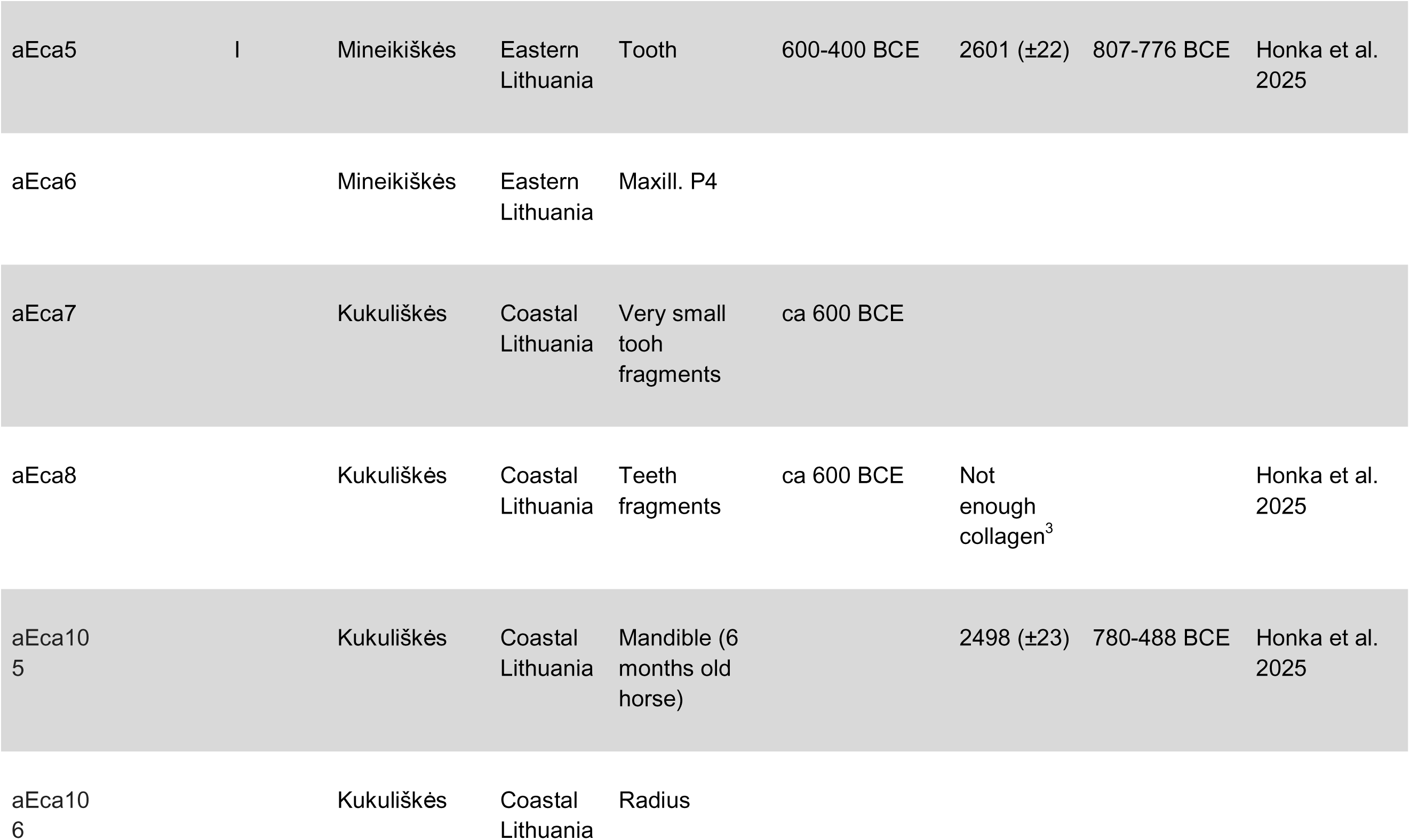
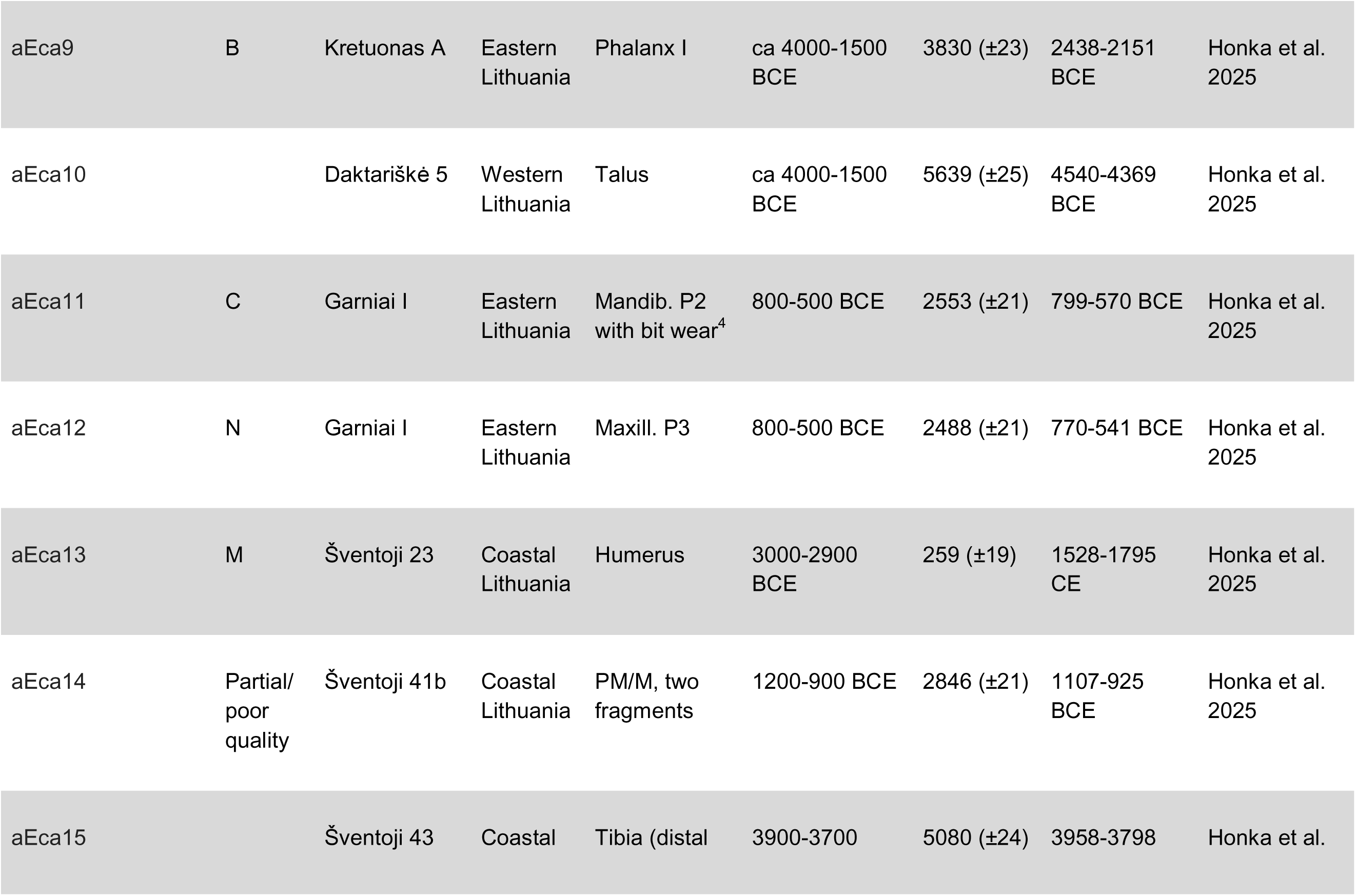
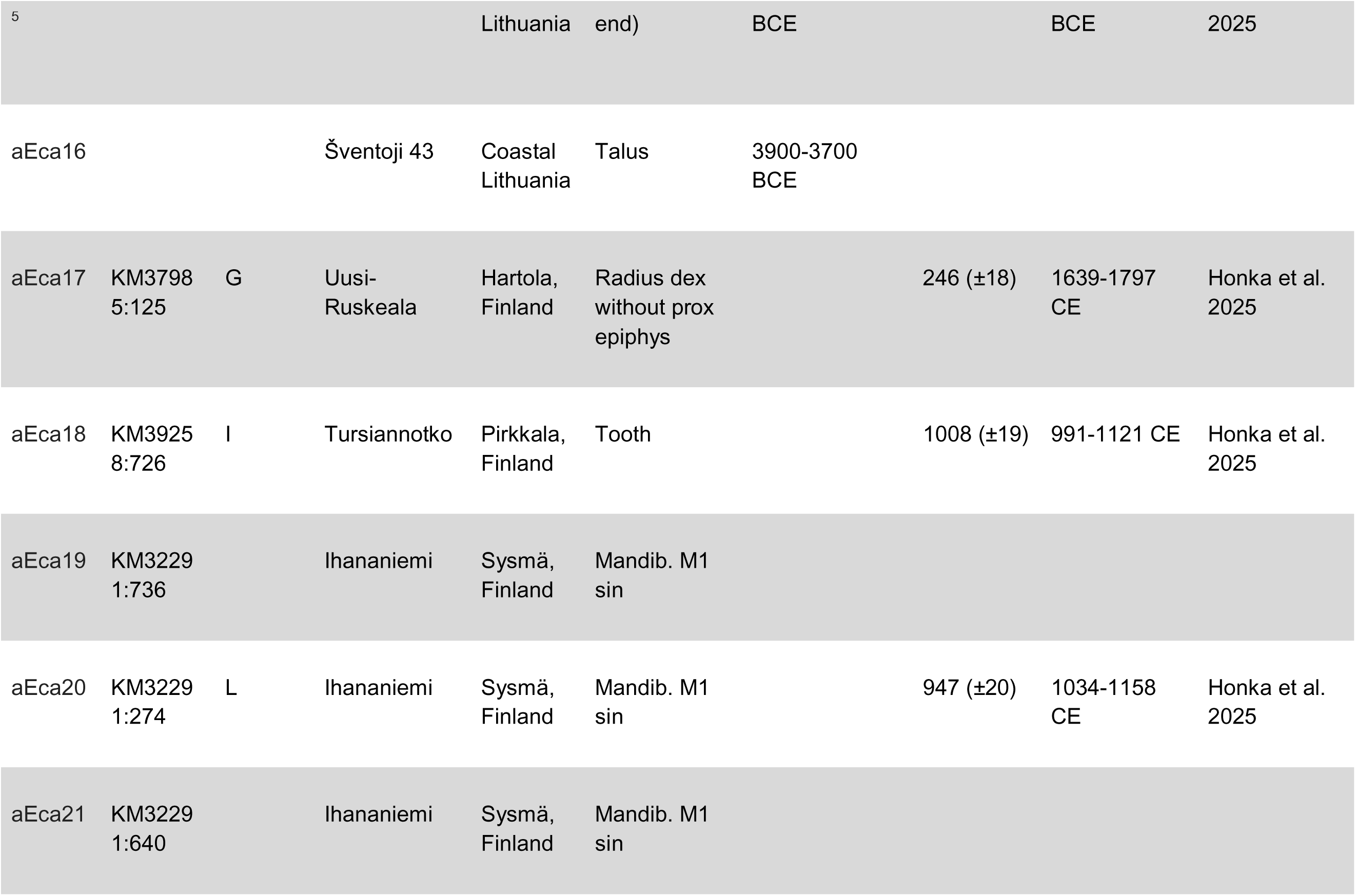
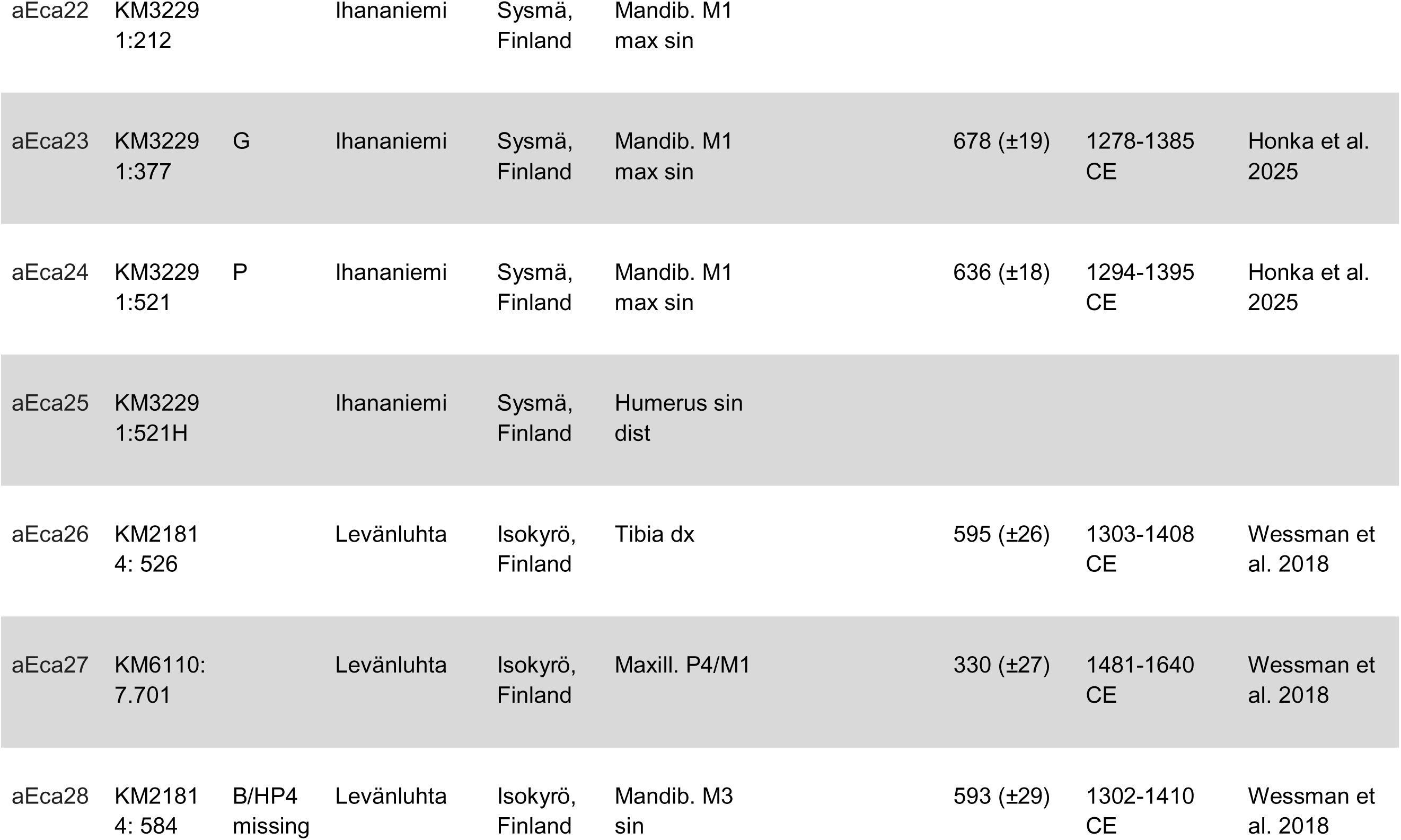
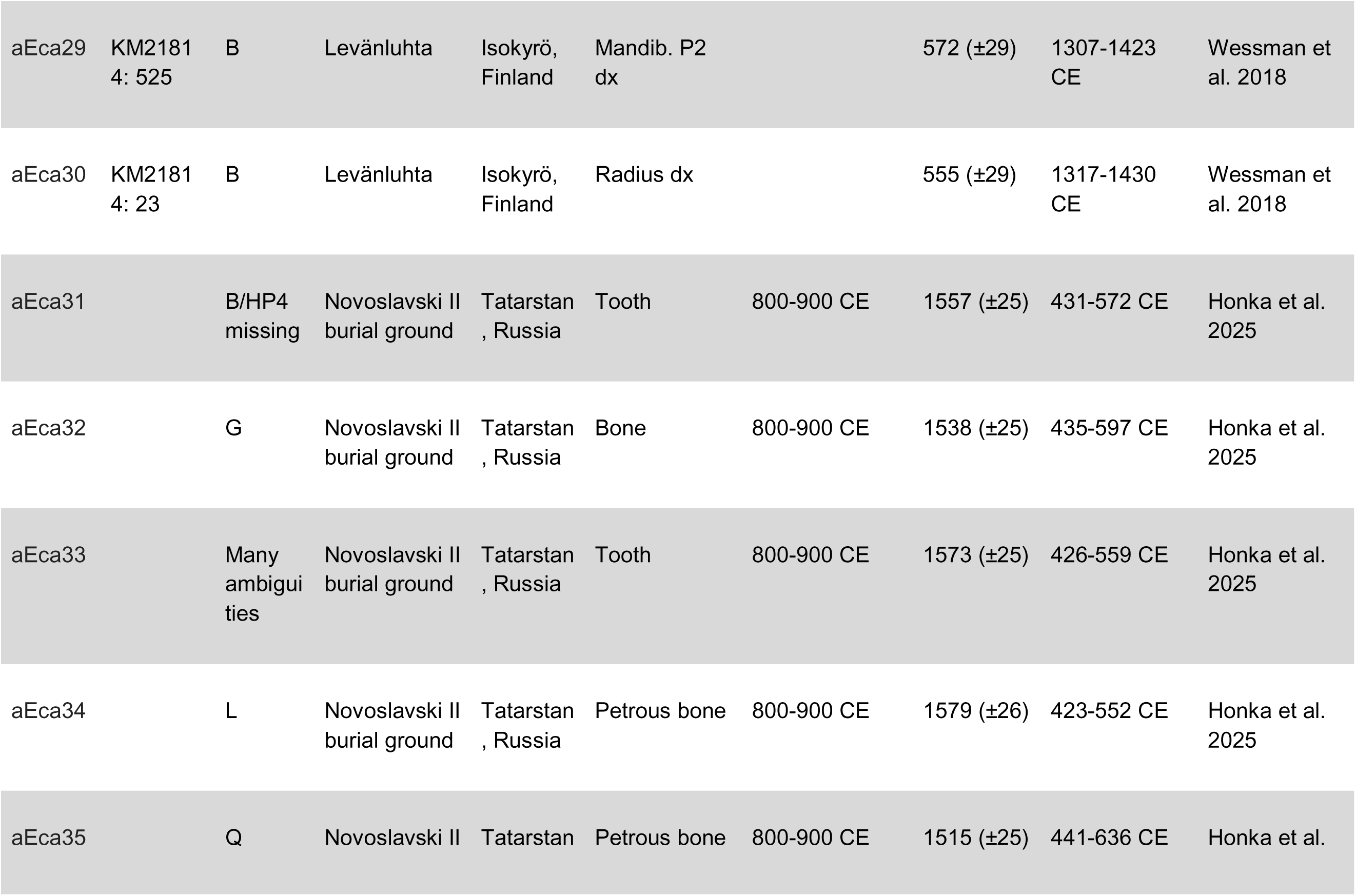
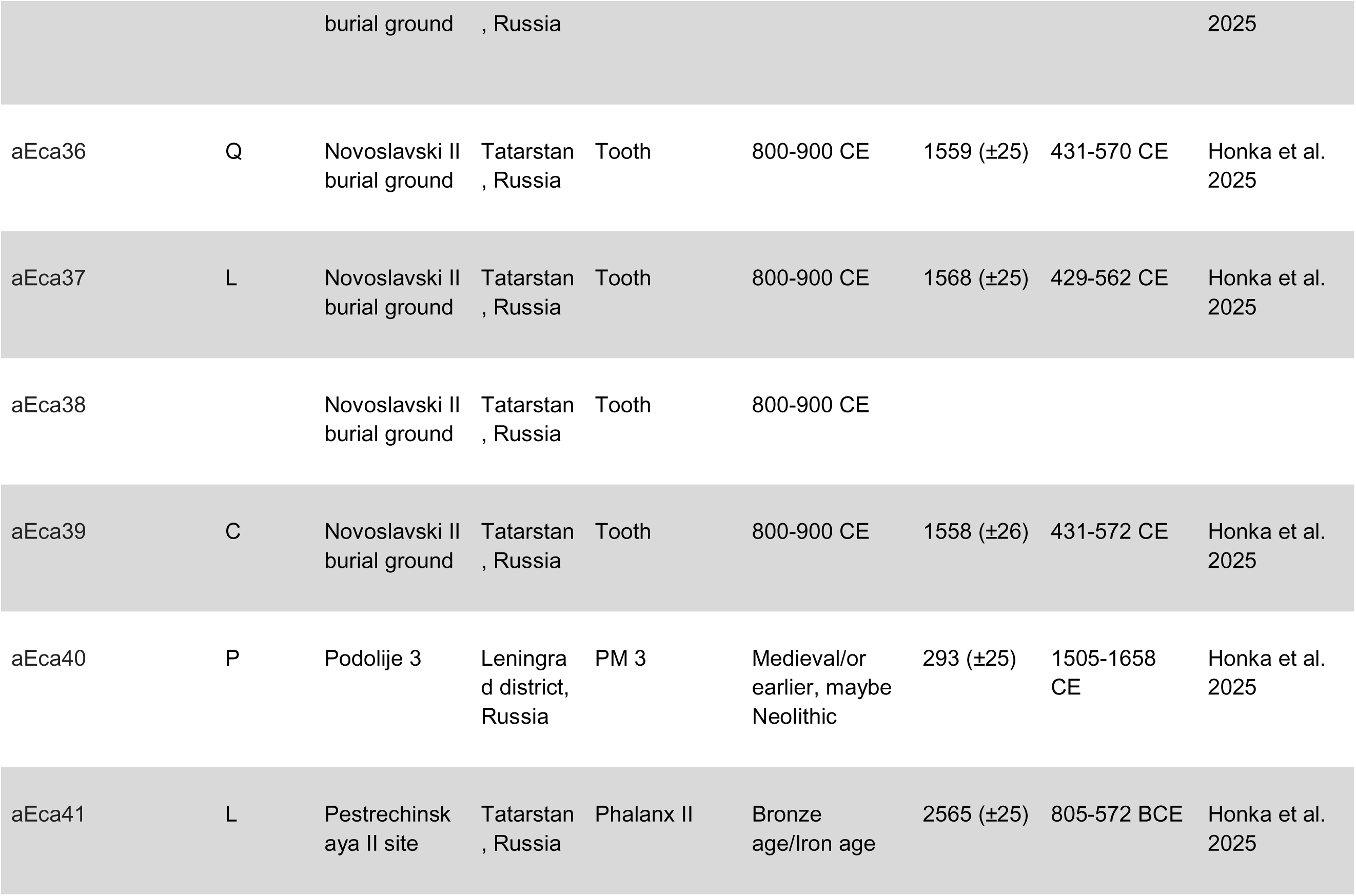
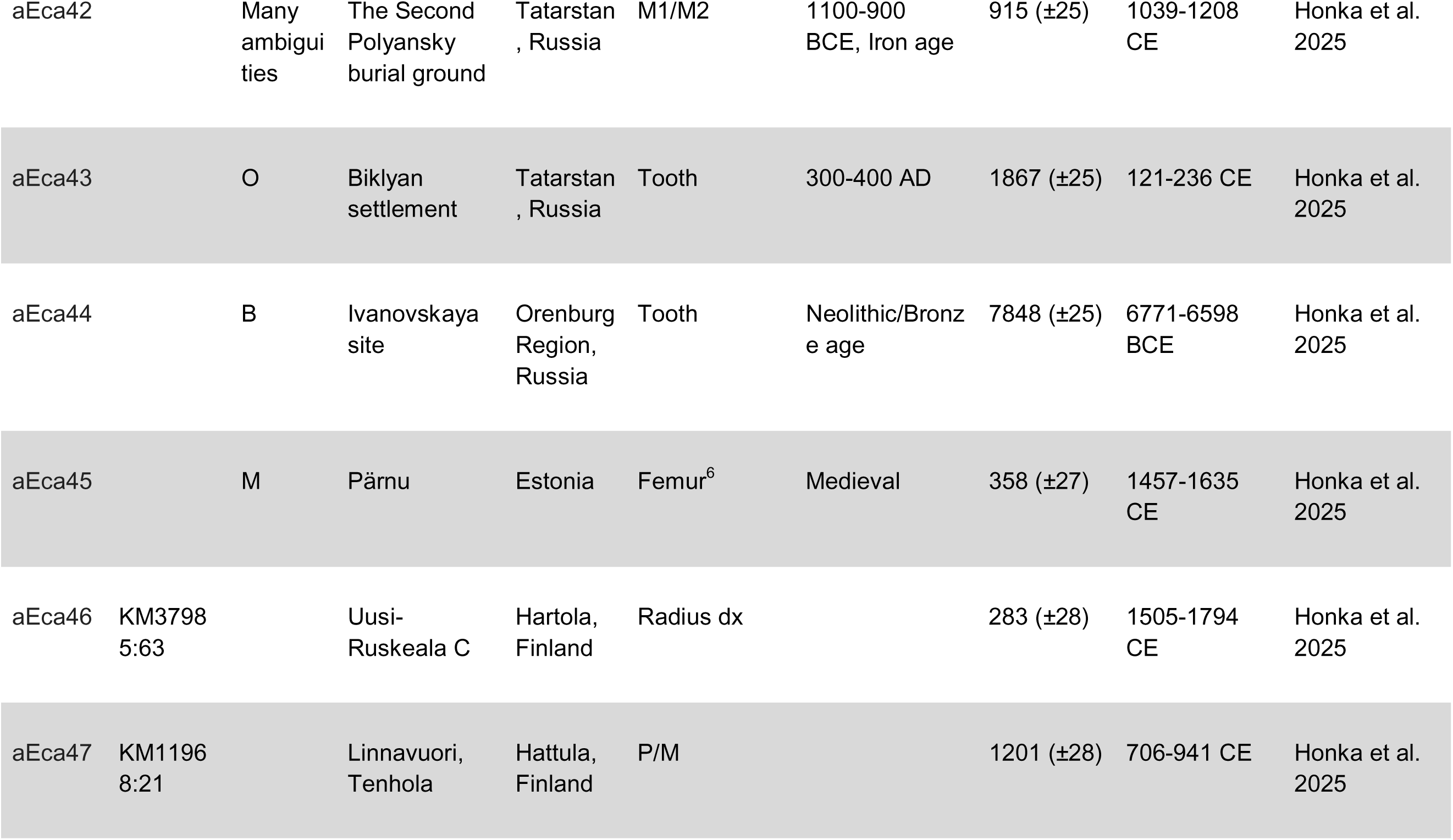
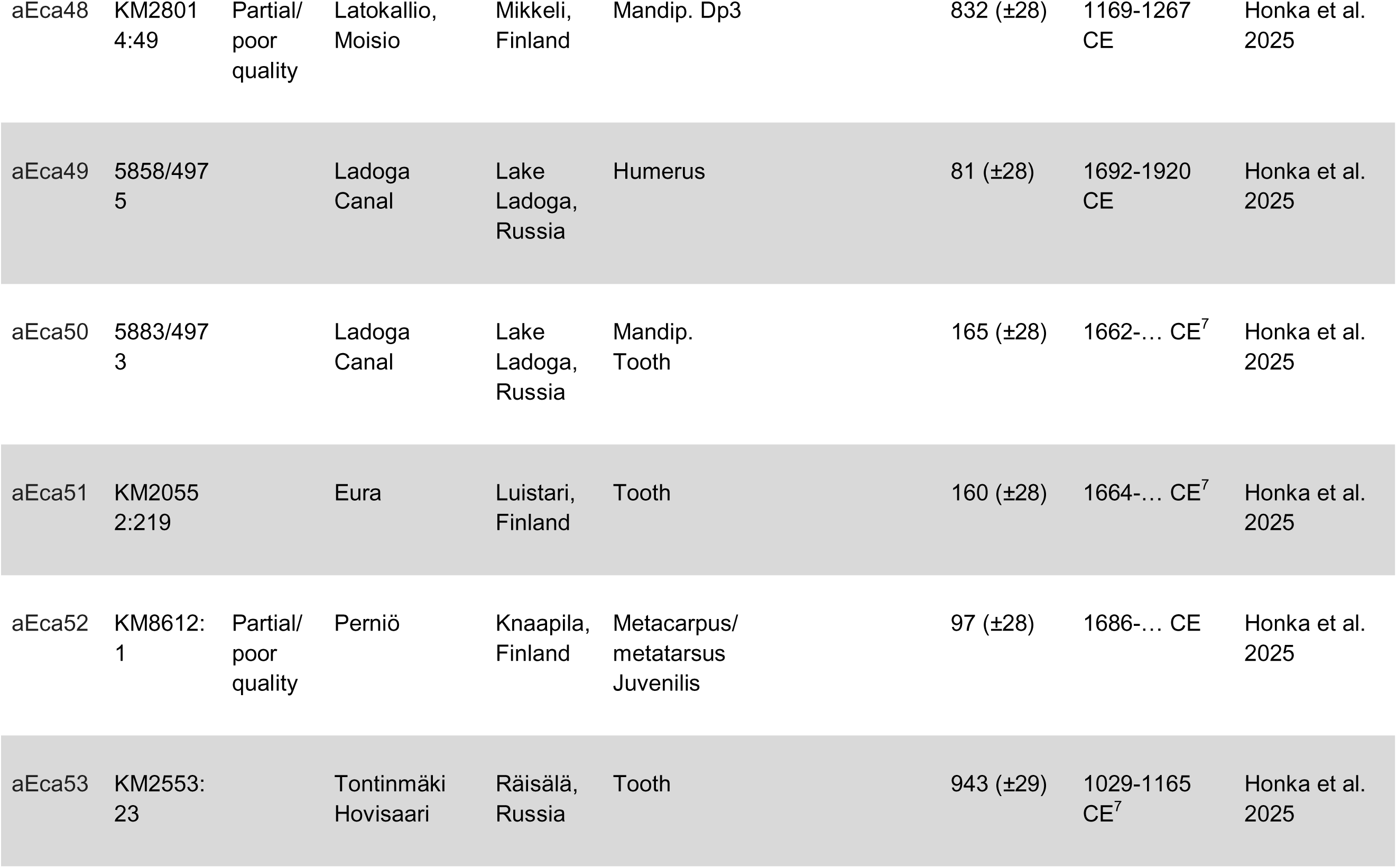
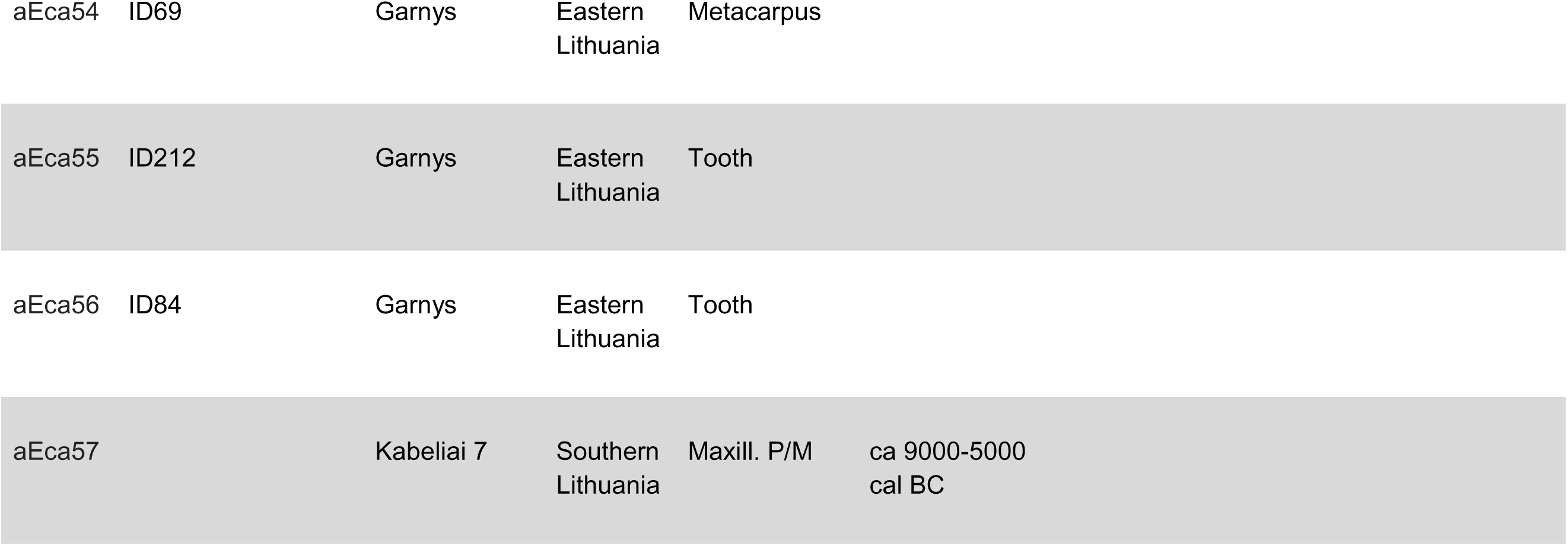
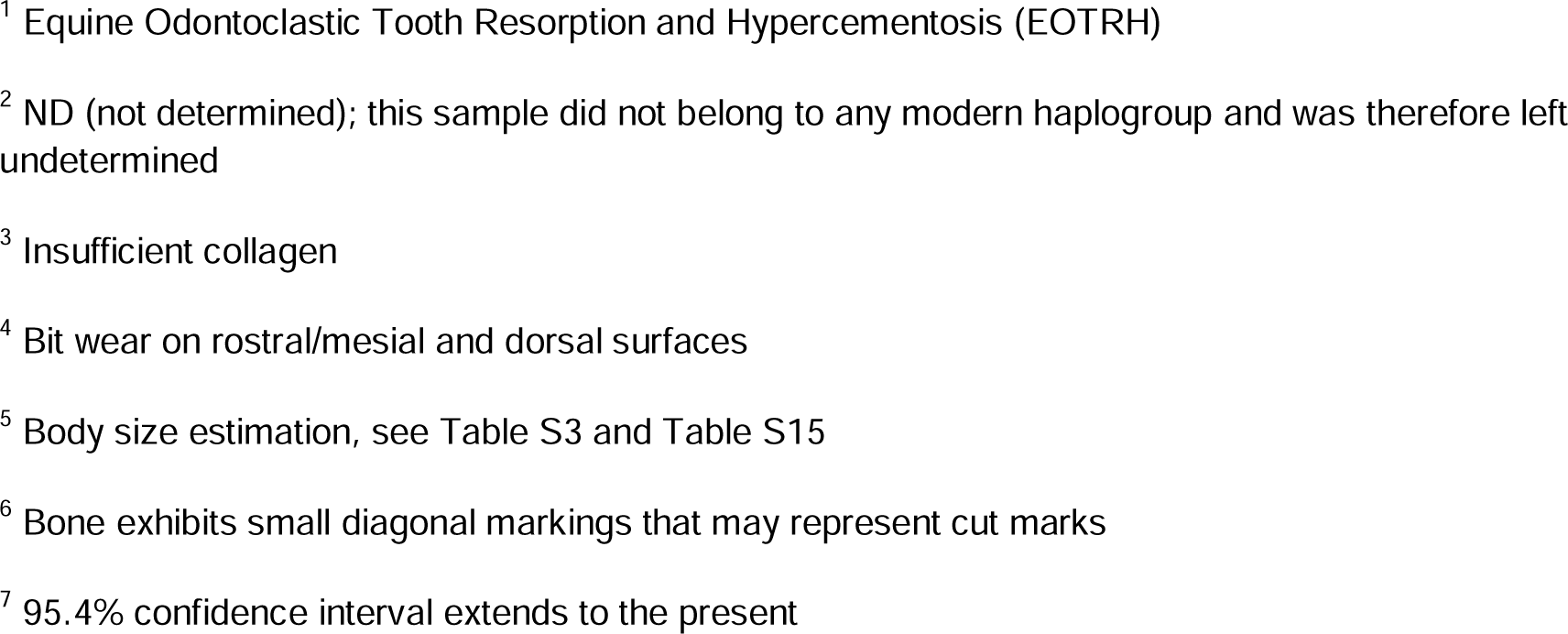
Archaeological bones and teeth of horses (*n* = 59) analysed for ancient DNA. Sample name, museum code, mitochondrial (mtDNA) haplogroup based on naming by Achilli et al. (2012), archaeological site, location, studied skeletal element and dating based on archaeological context are indicated. Radiocarbon age (^14^C age yr BP) and calibrated radiocarbon age (Cal BCE/CE (95.4%), calibrated using Oxcal 4.4 IntCal20, based on previous publications, are also indicated. M = molar, PM = premolar.

### Pathologies

One tooth (sample aEca2) exhibited clear signs of dental pathology (inflammation), while another showed possible signs of bit wear (sample aEca11), as reported for the first time in Minkevičius et al. (2023). Both teeth were evaluated by equine veterinarians to confirm the observed pathologies.

### Samples for ancient DNA

For ancient DNA, a total of 59 samples were analysed, originating from Lithuania (*n* = 22), Finland (*n* = 19), Russia (*n* = 17) and Estonia (*n* = 1) (Table 1, Fig. 2). From each site, we tested the same skeletal element or individuals of different ages. Thirty-nine of the samples were radiocarbon dated and analysed for stable carbon and nitrogen isotopes in previous publications (Honka et al., 2025; Oinonen et al., 2020; Wessman et al., 2018). The archaeological contexts were described in Honka et al. (2025), except for Levänluhta in Finland, a water burial site (see Data S1). The Lithuanian samples included two individuals most likely representing late wild horse populations (4539–4367 cal BCE and 3956–3797 cal BCE), one horse from a probably even older context (9000-5000 cal BCE; Gaižauskas et al. 2022), but not directly dated, and one that could be either wild or domestic based on its dating (2436–2149 cal BCE) (Honka et al., 2025). One wild horse from the Russian Ivanovskaya site (6770–6597 cal BCE; Honka et al., 2025) was also included.

### Ancient DNA extraction and mitochondrial DNA amplification

We extracted aDNA from the 59 bones or teeth (Table 1) following the protocol in Honka et al. (2018), in the clean-room facilities dedicated to aDNA at the Centre for Material Analysis at the University of Oulu, Finland. Precautions to avoid contamination are also described in Honka et al. (2018), and negative controls were included in each step. We amplified and sequenced a 573 bp fragment of the mitochondrial control region using an overlapping primer set (primer set 2, HP1-HP7, see Table S11 for correct primer orientation) from Cieslak et al. (2010). PCR reaction conditions are detailed in Methods S2. We sequenced the amplicons in both directions using the PCR primers and the BigDye Terminator v.3.1 (Applied Biosystems) chemistry. Sequencing reactions were run on an ABI 3730 (Applied Biosystems). Each fragment was amplified and sequenced at least twice (or more if necessary) to identify post-mortem changes or polymerase errors. CodonCode Aligner v.4.0.4. (CodonCode Corporation) was used for manual sequence editing and base-calling when necessary. We searched for identities between our sequences and those in GenBank using BLAST (basic local alignment tool).

BioEdit 7.2.5 (Hall, 1999) was used to align the obtained sequences with ancient horse sequences (73 sequences) and a set of modern horses divided into Finnhorse, eastern breeds and other horse groups (861 sequences), see Method S2 and Table S12. Additionally, mitochondrial genomes of 81 modern horses (GenBank accession numbers: JN398377–JN398457) were used to identify the 18 haplogroups defined by Achilli et al. (2012). Due to the large number of sequences, we used only distinct haplotypes identified with DnaSP v. 5.10 (Librado & Rozas, 2009). A median-joining haplotype network (Bandelt et al., 1999) was constructed using PopART (Leigh & Bryant, 2015). We constructed a separate haplotype network including two samples (aEca28 and aEca31) for which sequencing failed with primer pair HP4. We removed HP4 fragments from all sequences and concatenated the remaining two fragments (508-509 bp), allowing us to assign haplogroups to these two samples. These two samples were otherwise excluded from analyses. We constructed a maximum likelihood phylogenetic tree using the Tamura 3-parameter model (Tamura, 1992) with gamma distribution and invariant sites (TN92+G+I) with five discrete gamma categories, using all sites and 1000 bootstrap replications in MEGA 7.0.18 (Kumar et al., 2016). Otherwise, default settings were used for all parameters. The best nucleotide substitution model was also selected using MEGA. We also constructed a temporal statistical parsimony haplotype network using the TempNet (Prost & Anderson, 2011) R-script (R Core Team, 2023), based only on our ancient horses. Two samples (aEca3-4) were excluded from this analysis due to a lack of ^14^C dates.

Further, we created a presence-absence seriation matrix of the haplogroups for Lithuania, Finland and the Middle Volga region (representing a west-east geographic gradient) using PAST v. 4.12 (Hammer et al., 2001) to visualize haplogroup sharing across regions. Due to the limited sample size, we estimated genetic diversity for each country by including samples from all time periods, except for Estonia (only one sample), and excluding the single horse older than 3000 cal BCE with amplifiable DNA (sample aEca44). The number of haplotypes (*H*), haplotype diversity (*h*) and nucleotide diversity (π) were calculated using DnaSP v. 5.10 (Librado & Rozas, 2009). Ambiguous bases in the data were resolved according to the consensus derived from modern sequences, as the ambiguities occurred at conserved sites.

## Results

### Body size estimations

The Ivanovskaya site horses averaged 141.5 cm tall at the withers and 141.1 cm at the croup when adjusted for *E. caballus* (Table S13). These averages are 141.3 cm and 142.1 cm, respectively, when adjusted for *E. przewalskii.* Their average body mass was 410.3 kg. (Table S13). We adjusted for both subspecies, as it is unclear which provides a more appropriate reference. Mean withers heights of horses associated with Sredny Stog, Botai and Tersek cultures ranged between 139.3 cm (*E. caballus* adjusted) or 135.3 cm (*E. przewalskii* adjusted) to 140.7 cm (*E. caballus* adjusted) or 136.6 cm (*E. przewalskii* adjusted), whereas body masses ranged from 364.4 to 384.4 kg (Table S14). Body mass of the Lithuanian wild horse (aEca15) was 265.5 kg (Table S15).

The two Lithuanian Late Bronze Age phalanxes from Mineikiškės were from small horses with withers height of ∼119.3 cm, croup height of ∼120.8 cm and body weight of 275.8 kg (Table S15). It is possible that the phalanxes are actually from the same individual, but based on 95% confidence intervals, these likely originate from different individuals. The Lithuanian Pre-Viking Age horses (∼100-700 CE) averaged 127.4 cm at the withers (121.8-131.8 cm) with an average body mass of 306.6 kg (249.5-383.0 kg) (Table S16). Viking Age horses (∼800-1100 CE) were slightly smaller, averaging 124.5 cm at the withers (117.1-134.6 cm) with an average body mass of 276.3 kg (222.7-374.0 kg) (Table S17). Medieval horses (∼1200-1400 CE) averaged 125.0 cm at the withers (108.2-142.5 cm) and had an average body mass of 269.6 kg (172.5-385 kg) (Table S18). Their maximum rider weights (XRWs) were 92.1 kg, 85.8 kg and 79.7 kg, respectively (Tables S16-S18). Withers height estimations introduced here differ somewhat from those based on May (1985) (Table S19-S21). For horse aEca4, we were only able to estimate body mass, and this was 310.2 kg if the phalanx I (XX) was from the forelimb and 303.9 kg if it was from the hindlimb. For the Late Bronze-Early Roman Antilgė distal tibia, the body mass estimate was 200.4 kg (Table S15).

### Pathologies

One tooth from a Lithuanian horse (aEca2) showing signs of inflammation (Fig. 3A) was diagnosed as Equine Odontoclastic Tooth Resorption and Hypercementosis (EOTRH; Staszyk et al., 2008) by equine veterinarians. Another tooth from a Lithuanian horse (aEca11) exhibited signs of bit wear on the rostral/mesial and dorsal surfaces (Fig. 3B), as diagnosed by an equine veterinarian specialising in equine dentition. This horse was dated to 798–568 cal BCE (95.4% probability).

**Fig. 3.**
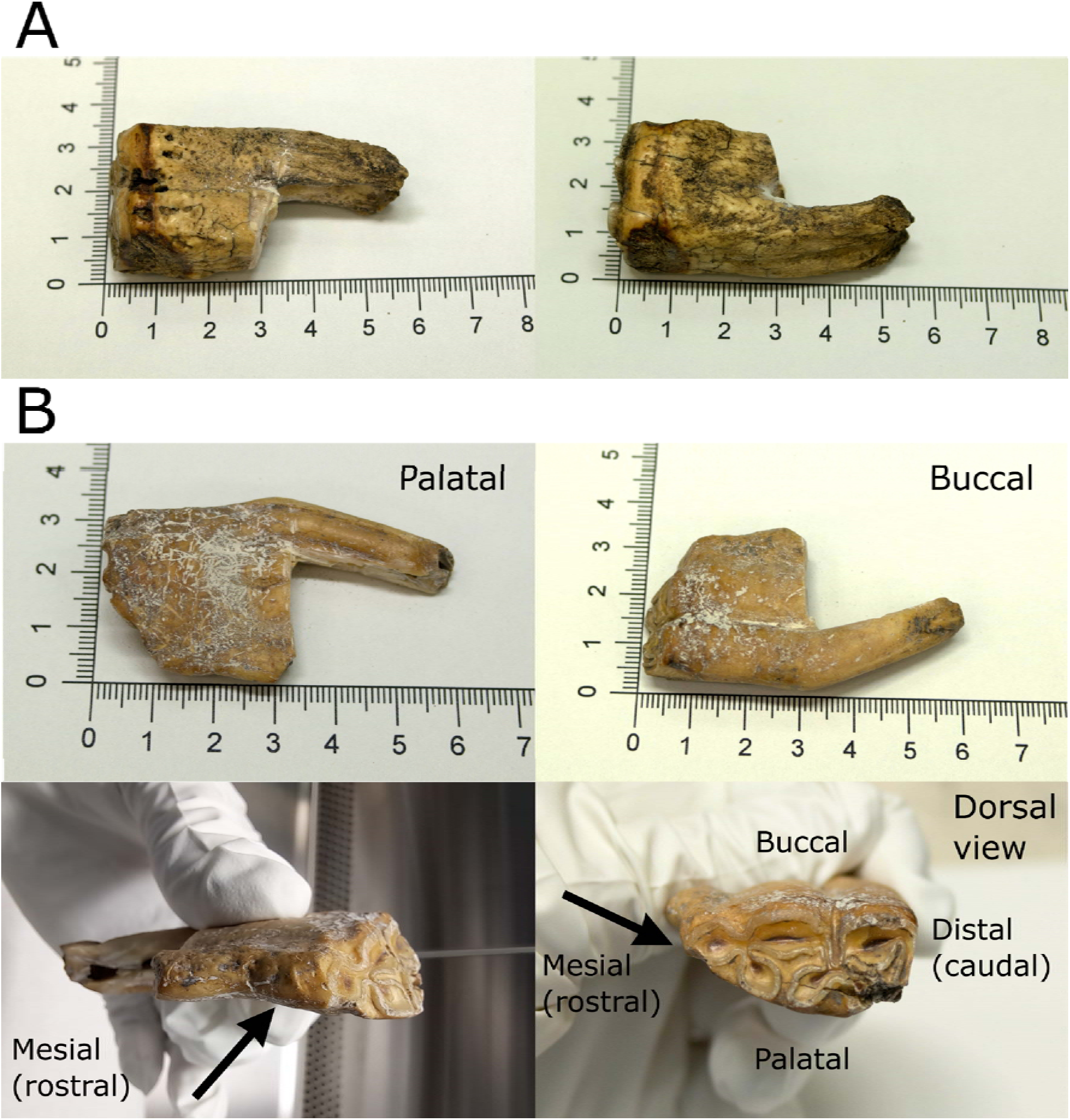
Two teeth with pathologies. **A)** Lithuanian horse tooth diagnosed with EOTRH (Equine Odontoclastic Tooth Resorption and Hypercementosis). The other root of the tooth was removed for ancient DNA analysis, radiocarbon dating and stable isotope analyses. **B)** Lithuanian horse tooth (lower P2) showing signs of bit wear in rostral/mesial and dorsal surfaces, indicated by arrows. The other root of the tooth was removed for ancient DNA analysis, radiocarbon dating and stable isotope analyses.

### Ancient DNA

Thirty-four out of 59 samples (58%) yielded amplifiable DNA (Table 1). However, four of these showed poor amplification with several primer pairs, and two showed numerous ambiguous bases; these were excluded from the analyses. Sequencing with primer pair HP4 failed for two samples; these were also excluded from the analyses, except for haplogroup assignment via a haplotype network (see methods). This resulted in 26 samples with 572–573 bp sequences (including one indel) for analyses (44%; Table 1). In seven samples, one or two primer pairs failed to amplify in the second or third repetition, but these were still included in the sequence analyses. Ten samples were successfully sequenced from Russia (59% success rate, total *n* = 17), eight from Lithuania (36% success rate, total *n* = 22), seven from Finland (37%, total *n* = 19) and one from Estonia (100%, total *n* = 1). Post-mortem changes or polymerase errors in PCR amplification were detected in some samples (nucleotide differences between two independent PCR-replicates). In these cases, we re-amplified and sequenced the fragments a third or fourth time. If nucleotides remained unresolved despite multiple attempts, we used IUPAC ambiguity codes.

Our ancient horses belonged to eleven of the eighteen haplogroups described by Achilli et al. (2012), missing haplogroups A, E, F, H, J, K and R (Fig. 4). The Russian samples belonged to haplogroups B, C, G, L, O, P and Q. The oldest horse from Russia (aEca44) belonged to haplogroup B and did not share its haplotype with modern or ancient reference horses, nor with any other horse in GenBank according to a BLAST search (Fig. 4, Fig. S1). The Finnish samples belonged to haplogroups B, G, I, L and P (Fig. 4). Lithuanian samples belonged to haplogroups B, C, D, I, M and N, but for one sample (aEca4), a haplogroup could not be assigned (Fig. 4), as it did not cluster with the defined haplogroups. Similar results were obtained from the maximum likelihood tree and the haplotype network analysis (Fig. S2). None of the oldest Lithuanian horses yielded amplifiable DNA despite several attempts, including trials from another bone of the same individual. The Lithuanian sample dated to ∼2200 BCE belonged to haplogroup B. The single Estonian horse (aEca45) in our dataset belonged to haplogroup M and shared its haplotype with a Lithuanian horse (aEca13; Fig. 7A). Two Finnish samples from the same site (aEca29 and aEca30; Levänluhta, Isokyrö) and with similar ages (Fig. 5A) shared the same haplotype (haplogroup B), as did a third individual from the same site and age based on a shorter concatenated sequence. No clear patterns were evident from the haplotype network that also included modern breeds from the studied region (Fig. S1). Based on the seriation matrix, only haplogroup B was shared among all geographic regions: Lithuania, Finland and the Middle Volga region (Fig. 5B). It was also the only haplotype shared between two temporal periods (Russian samples aEca34 and aEca41; Fig. 5A). Haplotype (*h*) and nucleotide (π) diversities were high, as almost no haplotypes were shared (Table 2).

**Fig. 4.**
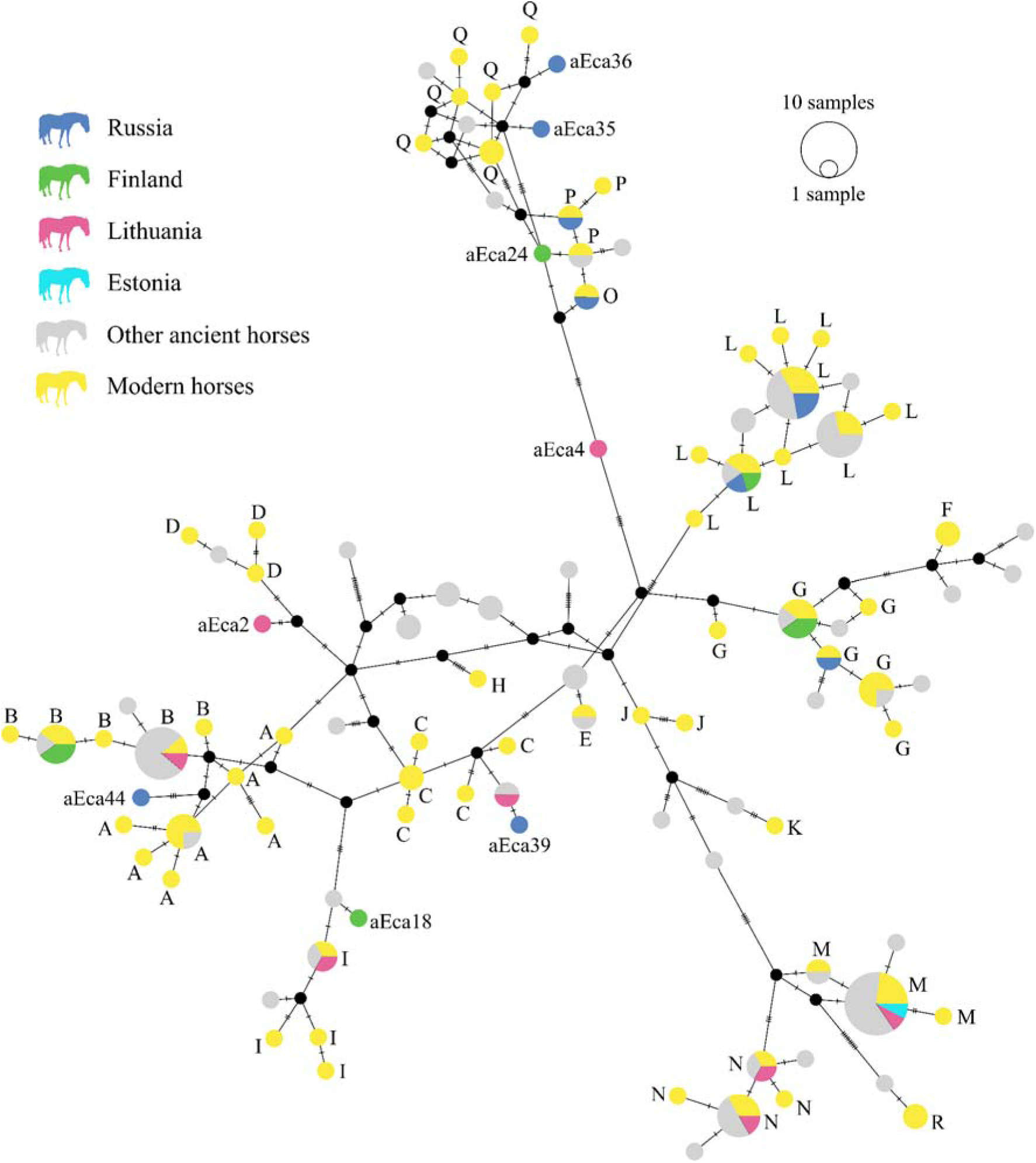
A median-joining haplotype network of 572-573 bp from the mitochondrial control region of subfossil horses, including one wild horse (aEca44), ancient horses downloaded from GenBank, and 81 sequences from Achilli et al. (2012) used as references to determine the major haplogroups. Different countries of origin are shown with different colours. The size of each circle is proportional to the frequency of each haplotype, and tick marks across branches indicate mutational differences.

**Fig. 5.**
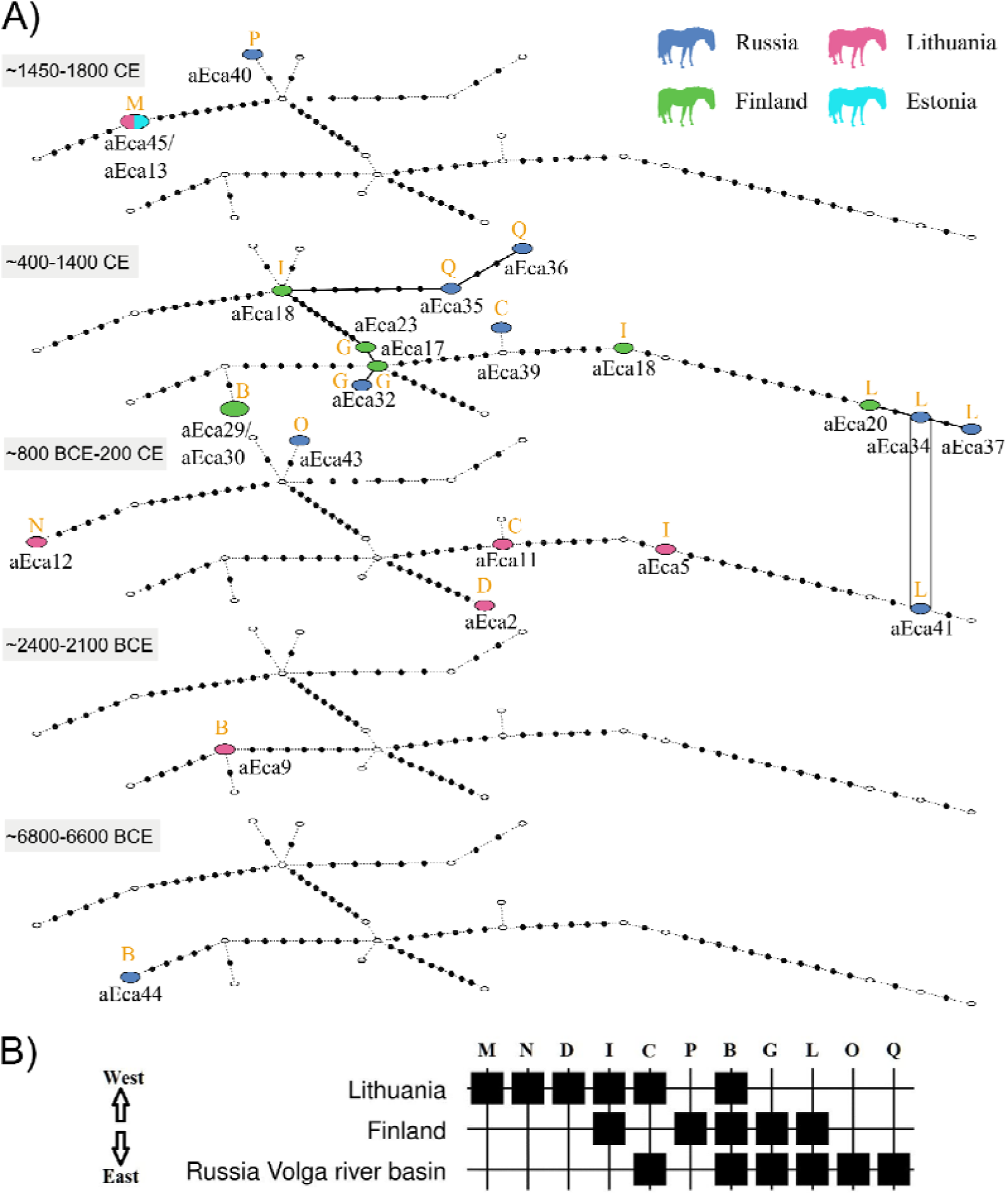
Temporal and geographic haplotype sharing among subfossil horses and one wild horse (aEca44). A) Temporal statistical parsimony network of the 572-573 bp from the mitochondrial control region. The size of each ellipse is proportional to the frequency of each haplotype, with the smallest representing one haplotype and the others representing two haplotypes. Countries of origin are shown in different colours, and sample names are provided. Small white ellipses denote haplotypes missing from that period, and small black dots indicate mutational differences between haplotypes. Note that the periods are discontinuous. Haplogroups, based on Achilli et al. (2012), are also indicated as determined in Fig. 4. B). Matrix of ancient horse haplogroup distribution based on the presence and absence (seriation algorithm) along a geographic gradient (west–east) from Lithuania, Finland and the Middle Volga region.

**Table 2.** Summary statistics for the 572-573 bp fragment of the mitochondrial control region of subfossil horses in each studied country, excluding Estonia and the oldest horse from Russia. Shown are sample size (*n*), number of haplotypes (*H*), haplotype diversity (*h*), and nucleotide diversity (π) with standard deviations (SD).

| Country | $n$ | $H$ | $h$ (SD) | $\pi$ (SD) |
| --- | --- | --- | --- | --- |
| Russia | 9 | 8 | 0.972<br>(0.064) | 0.02316<br>(0.0019) |
| Lithuania | 8 | 8 | 1.000<br>(0.063) | 0.01936<br>(0.0017) |
| Finland | 7 | 6 | 0.952<br>(0.096) | 0.02081<br>(0.0031) |

## Discussion

We studied wild, possibly wild and domestic horses from the eastern Baltic Sea region and western Russia. We inferred the body sizes of the wild ancestors of DOM2 horses and examined temporal changes in body sizes in Lithuanian horses using updated equations. Additionally, we verified the oldest domestic horse in Lithuania (798 - 568 cal BCE) based on bit wear in one tooth. Further, we identified EOTRH (Equine Odontoclastic Tooth Resorption and Hypercementosis) in one tooth. Based on sequencing of the mitochondrial control region from 26 ancient samples, we identified multiple maternal lineages but found no clear spatial or temporal patterns in mtDNA haplogroups, which is not surprising given the large variation in mitochondrial DNA among modern horses. The wild horse from the Ivanovskaya site harboured haplogroup B, whereas we were unable to amplify any mtDNA from the oldest (wild) horses from Lithuania. Haplogroup B appeared to have been widespread, as it was recorded in Finland, the Baltic states and western Russia, and in the Ivanovskaya wild horse; however, this may reflect chance due to the small sample size.

### Body size estimations

We estimated that the mean withers height of 25 wild horses from the Russian Neolithic Ivanovskaya site was 141.3-141.5 cm (depending on whether *E. caballus* or *E. przewalskii* was used for adjustment), and the mean body mass was 410.3 kg (Table S13). These Neolithic horses from the easternmost parts of the Pontic-Caspian Steppe averaged somewhat taller and heavier than more recent horses from the western Pontic-Caspian Steppe, represented by the Eneolithic horses of the Stredni Stog culture from Deriivka (Ukraine), as well as horses of the Eneolithic Tersek and Botai cultures of the northern Kazakh steppe. The Ivanovskaya wild horses, most likely belonging to a genetic cluster named NEO-NCAS (Neolithic Northern-Caspian) by Librado et al. (2021), are part of a lineage leading to DOM2 horses. Horses in this area represent the first known steppe horses with substantial ancestry that later became prominent in C-PONT horses and reached their maximum in DOM2 horses (Librado et al. 2021; 2024). Further genomic work is warranted to confirm the NEO-NCAS ancestry. The Ivanovskaya wild horses were relatively large in size and therefore provided a significant amount of meat for human societies that hunted them. Wild horses were one of the most important game animals for hunter-gatherers and early herders, who lived between the lower Volga and Ural rivers (Vybornov et al., 2018: Table 1). However, it should be noted that body mass estimations are not as reliable as withers height estimations, because adult body mass fluctuates seasonally, whereas joint sizes remain fixed (except in cases of pathological changes) (Niskanen & Ruff, 2018). In addition, there is individual variation in the relationship between soft tissue mass and skeletal size. Furthermore, due to factors such as poor nutrition or other environmental conditions, horses might not have reached their theoretical sizes and weights. Interestingly, the single wild horse from Lithuania, from which we were able to estimate body weight, was considerably smaller than the Russian wild horses, weighing only 265.5 kg. This may point to regional differences or diminishing body sizes in late wild horse populations.

The two Lithuanian Late Bronze Age Mineikiškės horses (though possibly the same individual) were about 120 cm in withers height, weighed 250-300 kg (Table S15), and thus were small ponies. However, their domestic/wild status is uncertain. The Late Bronze-Early Roman Age Antilgė specimen weighed only ∼200 kg (Table S15) and was thus probably a very small horse, not suitable for riding by average-sized adults. Later Lithuanian, probably domestic, horses were, on average, 127.4 cm high at the withers before the Viking Age, 124.5 cm during the Viking Age, and 125.0 cm in the Medieval period. These horses were slightly shorter than prehistoric and historic (850 BCE-1800 CE) Estonian horses which averaged 132 cm (Rannamäe et al., 2025). Using the formula applied in this study, the estimated height of Lithuanian horses is approximately 2–3 cm lower than that obtained using the method proposed by May (1985) in earlier research (Piličiauskienė et al. 2022). This difference, however, does not alter the main conclusions. In a previous analysis of 206 individuals (Piličiauskienė et al. 2022), the authors found that Lithuanian horses were generally shorter than those from Central and Western Europe and Scandinavia (Rosengren et al. 2017; Ameen et al. 2021; Chrószcz et al., 2021; Benkert, 2023). They also showed that horses in the pre-Viking period were larger than those from the Viking and medieval periods. Moreover, an important shift was observed in the Middle Ages, when horse size became far more variable than in earlier periods, and large individuals measuring 140–155 cm began to appear. These developments have been interpreted as the result of broader historical processes, including the arrival of the Teutonic Order in the eastern Baltic, intensified Lithuanian military campaigns, and growing contacts with both the German orders and Eastern Slavic lands (Piličiauskienė et al., 2022). The Lithuanian horses were about the size of modern Gotland Russ ponies and likely had a similar overall build. This has implications for their weight-carrying capacity and their suitability for riding and use as packhorses. Their average maximum rider weights (XRWs), ranging from 80-92 kg, are consistent with expectations for horses of this size, although variation exists in these values. Some horses could have carried rider weight well over 100 kg based on their metapodials, making some individuals suitable as mounts for knights and other armoured men-at-arms. The average sizes of Pre-Viking and Viking period males in Lithuania were, on average, 174.1 cm (Pre-Viking Plinkaigalis cemetery), 170.5 cm (Viking age Marvelė cemetery), and 172.9 cm and 75-77 kg (Pre-Viking Obeliai cemetery) (Jankauskas, 2001), thus capable of riding the small Lithuanian horses. However, the true weight-carrying ability of a horse is determined by its weakest point, whether insufficient size or muscle strength, or a conformation flaw such as a weak back or incorrect limb posture (Niskanen 2023: Supplementary information B). Estonian archaeological horses were calculated to have maximum rider weights of 86 kg, on average (Rannamäe et al., 2025), thus in the same range as the Lithuanian horses.

### Pathologies

We observed clear pathological changes in two teeth, as confirmed by equine veterinarians. One horse from Lithuania (aEca2; 18-202 cal CE) suffered from a painful EOTRH (Equine Odontoclastic Tooth Resorption and Hypercementosis) dental condition, which primarily affects the canine and incisor teeth, sometimes also the molars, and is characterized by resorptive or proliferative changes in the calcified dental tissues typically seen in older horses (> 15 years). Thus, this tooth likely belonged to a horse of advanced age. We are not aware of this condition being reported in ancient horses, although abnormal tooth wear and inflammation of tooth roots have been documented, for example, by Piličiauskienė et al. (2022).

Another tooth from Lithuania (aEca11) showed clear signs of bit wear on the rostral/mesial and dorsal surfaces of the lower P2 tooth, as observed in Minkevičius et al. (2023). Bit wear appears as a vertical strip of wear through the cementum exposing enamel (and in severe cases, dentine) on the mesial edge of the P2 tooth, the tooth that comes in contact with the bit. This specimen was dated to 798 - 568 cal BCE. The first evidence of bit wear on horse teeth comes from the Botai culture 3500-2700 BCE (Brown & Anthony, 1998; Outram et al., 2009), although some scholars interpret this as natural damage to teeth (Taylor & Barrón-Ortiz, 2021). This observation of bit wear is the oldest in the Baltic countries (Minkevičius et al. 2023) and thus is the oldest domestic horse in Lithuania (^14^C-dated in Honka et al. 2025). This indicates that domestic horses were transported to Baltic states by at least 800-570 BCE. However, cheek-pieces of bridles are known in the same time frame from Estonia (1100-500 BCE) (Maldre & Luik, 2009). As we were able to obtain mtDNA from this specimen, this sample is an excellent candidate for further genomic analysis.

### Maternal lineages in ancient DNA

We obtained mtDNA from the oldest horse of our dataset, the Ivanovskaya wild horse (6771-6598 cal BCE; ^14^C date Honka et al., 2025), but we were unable to recover DNA from wild horses in Lithuania despite multiple attempts. We could not analyse the oldest Finnish horse for aDNA because it is a burnt bone (820-546 cal BCE, Bläuer & Kantanen 2013), and DNA is destroyed by burning. The Ivanovskaya wild horse belonged to haplogroup B, similar to a Lithuanian horse dating to the period of the DOM2 migration. It is possible that this haplogroup was widespread in Europe during the Mesolithic and Neolithic, although based only on one wild and one potentially wild horse, this may be coincidental. Because most mitochondrial DNA lineages of modern horses predated domestication, mtDNA cannot serve as a marker for distinguishing wild from domestic horses (Cieslak et al., 2010). Evidence suggests that a large number of mares were involved in domestication, as indicated by the high mitochondrial DNA diversity observed in both ancient and modern horses (Cieslak et al., 2010; Achilli et al., 2012; Lippold et al., 2011). This finding implies either widespread domestication of wild horses or introgression of wild horse genes into domestic populations. The unique haplotype of the wild Russian horse (aEca44) was not shared with any other horse in GenBank based on a BLAST search, and may represent an extinct maternal lineage. Further genomic studies are warranted to clarify genetic relationships and diversity in ancient horses, and to distinguish wild from domestic individuals.

We discovered eleven different mtDNA haplogroups among our ancient horses, out of the 18 described by Achilli et al. (2012) (Fig. 4). Only haplogroup B was shared among all studied regions, Lithuania, Finland and the Middle Volga region (Fig. 5B), and thus appears to have been a widespread haplogroup among ancient Northeastern horses. However, it should be noted that, given the small sample sizes, the haplogroups identified may not reflect true population frequencies. Beyond this, we did not observe any clear spatial or temporal trends in mtDNA haplotypes. Most of the haplotypes found were shared with the modern horses used as the references (Fig. S1), except for one Lithuanian horse (aEca4; Fig. 4), for which we could not assign a haplogroup, probably because this mtDNA lineage may now be extinct. Interestingly, we noted that one of the 10^th^ -13^th^ CE Polish horses (Popovic et al., 2024), one ancient Chinese horse (800 BCE; Cai et al., 2007) and one Late Pleistocene wild horse from North-East Siberia (Cieslak et al., 2010) used here as a reference, belonged to the F-haplogroup, which is typical for the Przewalski’s horse. This was not reported in the original publications, likely because we used a fragment over twice as long, whereas the trimmed fragments may have lacked sufficient resolution to identify the F-haplogroup. Thus, the F-haplogroup may have been more widespread in the past than previously assumed.

## Conclusions

We found that the body sizes of wild horses excavated from the Russian Ivanovskaya site were similar to those of other wild horses, but a single Lithuanian wild horse was considerably smaller. Additionally, Late-Bronze Age-Early Roman Age horses were very small. We confirmed that Lithuanian domestic horses were pony-sized using updated equations for size estimation. As a new result, we estimated that the Lithuanian horses were capable of carrying riders and riding gear weighing 80-92 kg, and thus capable of carrying an average contemporaneous Lithuanian male. We also detected dental pathologies caused by EOTRH, and based on the bit wear traces, we confirmed the earliest domestic horse (Late Bronze Age) in the eastern Baltic region. Due to the lack of clear mitochondrial haplotype patterning in horses, we were unable to draw conclusions beyond describing the maternal haplogroups and their distribution. However, the samples with successful mtDNA amplification represent good candidates for future genomic studies, as DNA preservation was demonstrated in these specimens.

## Statements and declarations

### Ethics approval and consent to participate

Not applicable

### Consent for publication

Not applicable

### Availability of data and materials

Sequences of the ancient horses have been submitted to GenBank under accession numbers: PV252653-PV252678.

### Competing interests

The authors declare no competing interests.

### Funding

Analyses and DS have been funded by the Finnish Cultural Foundation. JH has been funded by the Biodiverse Anthropocenes Research Programme of the University of Oulu, supported by the Research Council of Finland (PROFI6 funding). LK and KM were partially funded by the Research Council of Finland (grant #370831).

### Authors’ contributions

JH, IVA, JA and LK designed and conceptualized the study arising from and during the symposium “*Archaeogenomics, step by step to understanding the history of domestic animals in Europe*”, held in Bolgar, Tatarstan, in 2020. IVA, DNS, OVA, AOA, SShA, RRV, MN, KM, and GP performed the archaeological examinations of the samples and bone measurements. JH, DS, and SO performed the laboratory work. JH analysed the data. MN and NP performed the calculations for the horse size estimations, and MN was primarily responsible for the review of the history of horse domestication. JH wrote the first version of the manuscript. All authors commented on previous drafts of the manuscript. All authors read and approved the final manuscript.

## Acknowledgements

We thank Pekka Moilanen and Kai Metsäkoivu for their assistance with the Centre for Material Analysis (CMA) clean room-facilities. We thank the Finnish Heritage Agency for granting permission to analyse the archaeological samples. We are grateful to the Museum of Staraya Ladoga for allowing us to examine their archaeological samples. We also thank equine veterinarians Johanna Weckmann and Mirjami Anttila for diagnosing pathologies in horse teeth. We are grateful to Kati Salo and Elisa Rautioaho for providing analysis results and information on the horse from Knaapila. We also thank Eve Rannamäe for reading the manuscript and providing helpful comments. In addition, we thank the two anonymous reviewers for their helpful comments on an earlier version of the manuscript.

## Notes

### Competing Interest Statement

The authors have declared no competing interest.

