## Supplementary material for "Body size, dental pathology and maternal genetic diversity of ancient horses in the eastern Baltic Sea region and western Russia"

### Supplementary files

**Table S1.** Full names and abbreviations of variables. Note that the terms metacarpal and metatarsal refer to metacarpal III and metatarsal III, respectively.

| **Full name(s)** | **Abbreviations used by von den Driesch (1976)** | **Abbreviations used based Eisenmann (1986)** |
|---|---|---|
| Withers height | -- | WH |
| Croup height | -- | CH |
| Body mass | -- | BM |
| Metacarpal greatest length | GL of metacarpal | MC1 |
| Metacarpal III lateral length | LI of metacarpal | MC1 |
| Distal articular breadth of metacarpal | Bd of metacarpal | MC11 |
| Shaft breadth of metacarpal | SD of metacarpal | MC3 |
| Proximal breadth of metacarpal | Bp of metacarpal | MC5 |
| Least depth of the medial condyle of metacarpal | -- | MC13 |
| Metatarsal greatest length | GL of metatarsal | MT1 |
| Metatarsal lateral length | LI of me | MT2 |
| Shaft breadth of metatarsal | SD of metatarsal | MT3 |
| Proximal breadth of metatarsal | Bp of metatarsal | MT5 |
| Distal articular breadth of metatarsal | Bd of metatarsal | MT11 |
| Least depth of the medial condyle of metatarsal | -- | MT13 |
| Greatest length of first phalanx of forelimb | GL of first phalanx of forelimb | P1fore1 |

| | | |
|---|---|---|
| Proximal breadth of first phalanx of forelimb | Bp of first phalanx of forelimb | P1fore4 |
| Distal supra-articular breadth of first phalanx of forelimb | Bd of first phalanx of forelimb | P1fore6 |
| Greatest length of first phalanx of forelimb | GL of first phalanx of hindlimb | P1hind1 |
| Proximal breadth of first phalanx of hindlimb | Bp of first phalanx of hindlimb | P1hind4 |
| Distal supra-articular breadth of first phalanx of hindlimb | Bd of first phalanx of forelimb | P1hind6 |
| Greatest length of second phalanx of forelimb | GL of second phalanx of forelimb | P2fore1 |
| Proximal breadth of second phalanx of forelimb | Bp of second phalanx of forelimb | P2fore4 |
| Distal supra-articular breadth of second phalanx of forelimb | Bd of first phalanx of forelimb | P2fore6 |
| Greatest length of first phalanx of hindlimb | GL of first phalanx of hindlimb | P2hind1 |
| Proximal breadth of second phalanx of hindlimb | Bp of second phalanx of hindlimb | P2hind4 |
| Distal supra-articular breadth of second phalanx of hindlimb | Bd of second phalanx of hindlimb | P2hind6 |
| Distal breadth of tibia | Dd of tibia | TIB7 |
| Distal depth of tibia | Bd of tibia | TIB8 |
| Relative distal articular breadth of metacarpal | -- | MC11/MC1x100 |
| Relative distal articular breadth of metatarsal | -- | MT11/MT1x100 |
| Relative distal breadth of first phalanx of forelimb | -- | P1fore6/P1fore1x100 |

| | | |
|---|---|---|
| Relative distal breadth of first phalanx of hindlimb | -- | P1hind6/P1hind1x100 |
| Relative distal breadth of second phalanx of forelimb | -- | P2fore6/P2fore1x100 |
| Relative distal breadth of second phalanx of hindlimb | -- | P2hind6/P2hind1x100 |
| Metacarpal distal joint size | -- | MC11xMC13 |
| Metatarsal distal joint size | -- | MT11xMT13 |
| Tibial distal joint size | – | TIB7xTIB8 |

von den Driesch, A. 1976. A guide to the measurement of animal bones from archaeological sites: As developed by the Institut Für Palaeoanatomie, Domestikationsforschung und Geschichte Der Tiermedizin of the University of Munich. Harvard University Press, Harvard.

**Table S2**. Measurements of horse bones from Ivanovskaya. Each bone is represented by its greatest length (GL), smallest midshaft breadth (SD), greatest proximal breadth (Bp) and greatest distal breadth (Bd). See Table S1 for the full names of the abbreviations of variables. The measurements are in millimeters (mm).

| Specimen | Bone | MC1<br>(GL) | MC3<br>(SD) | MC5<br>(Bp) | MC11<br>(Bd) | MT1<br>(GL) | MT3<br>(SD) | MT5<br>(Bp) | MT11<br>(Bd) |
|---|---|---|---|---|---|---|---|---|---|
| 1 | Metacarpal III | 226.0 | 35.0 | 48.0 | 49.0 | | | | |
| 2 | Metacarpal III | 218.0 | 35.0 | 51.0 | 49.0 | | | | |
| 3 | Metatarsal III | | | | | 271.0 | 37.0 | 52.0 | 56.0 |
| 4 | Metatarsal III | | | | | 260.0 | 34.0 | 50.0 | 53.0 |
| | | **P1fore1<br>(GL)** | **P1fore3<br>(SD)** | **P1fore4<br>(Bp)** | **P1fore6<br>(Bd)** | **P1hind1<br>(GL)** | **P1hind3<br>(SD)** | **P1hind4<br>(Bp)** | **P1hind6<br>(Bd)** |
| 5 | Phalanx I (fore) | 88.0 | 39.0 | 58.0 | 49.0 | | | | |
| 6 | Phalanx I (fore) | 89.0 | 38.0 | 56.0 | 50.0 | | | | |
| 7 | Phalanx I (fore) | 91.0 | 41.0 | 61.0 | 52.0 | | | | |
| 8 | Phalanx I (fore) | 93.0 | 41.0 | 61.0 | 52.0 | | | | |
| 9 | Phalanx I (fore) | 87.0 | 39.0 | 56.0 | 51.0 | | | | |
| 10 | Phalanx I (fore) | 83.0 | 38.0 | 54.0 | 47.0 | | | | |
| 11 | Phalanx I (fore) | 81.0 | 35.0 | 55.0 | 44.0 | | | | |
| 12 | Phalanx I (fore) | 96.0 | 37.0 | 59.0 | 37.0 | | | | |
| 13 | Phalanx I (fore) | 92.0 | 36.0 | 58.0 | 36.0 | | | | |
| 14 | Phalanx I (fore) | 82.0 | 35.0 | 52.0 | 35.0 | | | | |
| 15 | Phalanx I (hind) | | | | | 92.0 | 37.0 | 55.0 | 48.0 |

| | | | | | | | | | |
|---|---|---|---|---|---|---|---|---|---|
| 16 | Phalanx I (hind) | | | | | 83.0 | 39.0 | 59.0 | 49.0 |
| 17 | Phalanx I (hind) | | | | | 76.0 | 35.0 | 54.0 | 46.0 |
| 18 | Phalanx I (hind) | | | | | 83.0 | 36.0 | 59.0 | 48.0 |
| 19 | Phalanx I (hind) | | | | | 89.0 | 36.0 | 54.0 | 44.0 |
| | | **P2fore1 or P2hind1 (GL)** | | **P2fore3 or P2hind3 (SD)** | | **P2fore4 or P2hind4 (Bp)** | | **P2fore6 or P2hind6 (Bd)** | |
| 20 | Phalanx II | 47.0 | | 46.0 | | 52.0 | | 46.0 | |
| 21 | Phalanx II | 49.0 | | 49.0 | | 55.0 | | 49.0 | |
| 22 | Phalanx II | 51.0 | | 44.0 | | 55.0 | | 44.0 | |
| 23 | Phalanx II | 53.0 | | 47.0 | | 58.0 | | 47.0 | |
| 24 | Phalanx II | 54.0 | | 47.0 | | 56.0 | | 47.0 | |
| 25 | Phalanx II | 52.0 | | 46.0 | | 54.0 | | 46.0 | |

**Table S3.** Sites, dates and measurements of horse bones from Lithuania predating the Common Era (CE). Each bone is represented by its greatest length (GL), smallest midshaft breadth (SD), greatest proximal breadth (Bp) and greatest distal breadth (Bd). See Table S1 for the full names of the abbreviations of variables. The measurements are in millimeters (mm). 14C-dating results are from Honka et al. 2025.

| **Site** | **Specimen** | **Date** | **P1fore1 (GL)** | **P1fore3 (SD)** | **P1fore4 (Bp)** | **P1fore6 (Bd)** | **P1hind1 (GL)** | **P1hind3 (SD)** | **P1hind4 (Bp)** | **P1hind6 (Bd)** | **TIB7 (Dd)** | **TIB8 (Bd)** |
|---|---|---|---|---|---|---|---|---|---|---|---|---|
| Šventoji 43 | aEca15* (Tibia) | 3881 BC ± 57 | -- | -- | -- | -- | -- | -- | -- | -- | 67.9 | 42.03 |
| Mineikiškės | Phalanx I (fore)** | Late Bronze Age | 71.2 | 30.2 | 50.6 | 42.5 | -- | -- | -- | -- | | |
| Mineikiškės | Phalanx I (hind)** | Late Bronze Age | -- | -- | -- | -- | 70.9 | 30.0 | 48.1 | 41.1 | | |
| Antilgė | Phalanx I (fore or hind?) | Late Bronze Age – Early Roman | 87.22 | | 51.17 | | 87.22 | | 51.17 | | | |
| Antilgė | Tibia | Late Bronze Age – Early Roman | -- | -- | -- | -- | -- | -- | -- | -- | 61.2 | 39.5 |

* See Table 1 in main text

** possibly the same individual

**Table S4**. Measurements of horse bones from Lithuania subdivided into three temporal samples. Each bone is represented by its greatest length (GL), smallest midshaft breadth (SD), greatest proximal breadth (Bp) and greatest distal breadth (Bd). Ll stands for lateral length (*lateral incisura*). See Table S1 for the full names of the abbreviations of variables. The measurements are in millimeters (mm). 14C-dating results are from Piličiauskienė et al. 2022.

| **Pre-Viking period Iron Age (*n* = 8)** | | | | | | | | | | |
|---|---|---|---|---|---|---|---|---|---|---|
| **Site** | **Specimen** | **Date** | **MC1 (GL)** | **MC2 (Ll)** | **MC11 (Bd)** | **MC13 (--)** | **MT1 (GL)** | **MT2 (Ll)** | **MT11 (Bd)** | **MT13 (--)** |
| Pagrybis cemetary | 142 | 6-7th c. CE | 210.0 | 203.5 | 47.5 | 27.9 | 252.0 | 246.0 | 47.7 | 26.9 |
| Pagrybis cemetary | 145 | 601–662 CE | 204.0 | 195.0 | 50.0 | 31.5 | -- | -- | -- | -- |
| Pagrybis cemetary | 157 | 540–640 CE | 206.0 | 197.0 | 45.2 | 27.6 | 246.0 | 237.0 | 42.7 | 25.1 |
| Pagrybis cemetary | 207 | 565–654 CE | 202.0 | 191.1 | 43.2 | 26.7 | 243.6 | 234.0 | 42.3 | 25.8 |
| Taurapilis barrow cemetery | 1 | 3-6th c. CE | -- | -- | -- | -- | 247.8 | 238.0 | 45.8 | 27.3 |
| Taurapilis barrow cemetery | 4 | 259–538 CE | 212.0 | 204.5 | 45.9 | 28.0 | 252.1 | 245.0 | 47.0 | 27.5 |
| Marvelė cemetery | 113 | 134–408 CE | 208.0 | 201.0 | 49.6 | 27.5 | 252.0 | 245.0 | 49.0 | 27.5 |
| Taurapilis barrow cemetery | 5 | 236–530 CE | 202.0 | 193.0 | 44.8 | 25.8 | -- | -- | -- | -- |
| **Viking period (*n* = 9)** | | | | | | | | | | |
| **Site** | **Specimen** | **Date** | **MC1 (GL)** | **MC2 (Ll)** | **MC11 (Bd)** | **MC13 (--)** | **MT1 (GL)** | **MT2 (Ll)** | **MT11 (Bd)** | **MT13 (--)** |
| Degsnė Labotiškė barrow cemetery | 5 | 9-11th c. CE | 195.0 | 186.0 | 44.8 | 25.6 | | | | |
| Marvelė cemetery | 35 | 9-11th.c. CE | 202.0 | 196.0 | 43.2 | 27.5 | | | | |
| Marvelė cemetery | 40 | 9-11th.c. CE | 220.0 | 210.3 | 49.4 | 31.6 | 255.0 | 247.5 | 49.2 | 30.3 |
| Marvelė cemetery | 49 | 9-11th.c. CE | 193.5 | 187.5 | 41.4 | 25.5 | | | | |
| Marvelė cemetery | 72 | 9-11th.c. CE | 208.0 | 201.5 | 45.0 | 27.3 | | | | |
| Marvelė cemetery | 103 | 9-11th.c. CE | 206.0 | 200.0 | 42.4 | 25.6 | 249.0 | 242.0 | 43.1 | 25.2 |
| Marvelė cemetery | 108 | 9-11th.c. CE | 210.0 | 204.0 | 46.3 | 27.0 | 260.0 | 247.5 | 45.9 | 25.2 |
| Marvelė cemetery | 116 | 9-11th.c. CE | 205.0 | 198.0 | 45.3 | 26.6 | 246.0 | 240.0 | 44.8 | 26.6 |
| Marvelė cemetery | 117 | 9-11th.c. CE | 192.0 | 190.0 | 42.4 | 26.4 | 236.0 | 230.0 | 44.1 | 26.5 |
| **Medieval period (*n* = 26)** | | | | | | | | | | |
| **Site** | **Specimen** | **Date** | **MC1 (GL)** | **MC2 (Ll)** | **MC11 (Bd)** | **MC13 (--)** | **MT1 (GL)** | **MT2 (Ll)** | **MT11 (Bd)** | **MT13 (--)** |
| Vilnius Lower castle | 67848 | 13-14th c. CE | 231.5 | 224.0 | 48.2 | 29.8 | | | | |
| Vilnius Lower castle | 14532 | 13-14th c. CE | 215.5 | 205.0 | 44.4 | 26.6 | | | | |
| Vilnius Lower castle | 66188 | 13-14th c. CE | 186.8 | 179.0 | 41.1 | 23.6 | | | | |

| | | | | | | | | | | |
|---|---|---|---|---|---|---|---|---|---|---|
| Vilnius Lower castle | 66189 | 13-14th c. CE | 193.0 | 185.5 | 42.4 | 23.1 | | | | |
| Vilnius Lower castle | 4796 | 13-14th c. CE | | | | | 218.0 | 211.0 | 40.0 | 22.8 |
| Vilnius Lower castle | 66191 | 13-14th c. CE | | | | | 240.8 | 234.0 | 42.0 | 24.5 |
| Kernavė medieval town | K24 | 13-14th c. CE | 209.0 | 202.0 | 44.1 | 24.8 | | | | |
| Kernavė medieval town | K30 | 13-14th c. CE | 233.0 | 225.0 | 50.8 | 30.0 | | | | |
| Kernavė medieval town | K31 | 13-14th c. CE | 229.0 | 222.0 | 48.4 | 30.1 | | | | |
| Kernavė medieval town | K32 | 13-14th c. CE | 224.0 | 216.0 | 46.1 | 26.2 | | | | |
| Kernavė medieval town | K33 | 13-14th c. CE | 221.0 | 211.0 | 47.8 | 27.1 | | | | |
| Kernavė medieval town | K35 | 13-14th c. CE | 205.0 | 197.0 | 46.2 | 27.0 | | | | |
| Kernavė medieval town | K36 | 13-14th c. CE | 178.0 | 169.0 | 38.6 | 23.3 | | | | |
| Kernavė medieval town | K37 | 13-14th c. CE | 217.0 | 209.0 | 48.2 | 27.6 | | | | |
| Kernavė medieval town | K38 | 13-14th c. CE | 203.5 | 194.0 | 45.6 | 26.2 | | | | |
| Kernavė medieval town | K39 | 13-14th c. CE | 198.0 | 193.0 | 43.1 | 25.6 | | | | |
| Kernavė medieval town | K40 | 13-14th c. CE | 183.0 | 175.8 | 40.6 | 24.2 | | | | |
| Kernavė medieval town | K41 | 13-14th c. CE | | | | | 271.5 | 265.0 | 48.0 | 29.2 |
| Kernavė medieval town | K42 | 13-14th c. | | | | | 261.0 | 251.0 | 45.2 | 26.2 |
| Kernavė medieval town | K43 | 13-14th c. | | | | | 250.0 | 251.0 | 41.7 | 24.4 |
| Kernavė medieval town | K44 | 13-14th c. | | | | | 241.0 | 237.0 | 44.2 | 25.4 |
| Kernavė medieval town | K45 | 13-14th c. | | | | | 237.0 | 229.0 | 45.7 | 26.6 |
| Kernavė medieval town | K45.1 | 13-14th c. | | | | | 244.0 | 234.0 | 41.6 | 24.6 |
| Kernavė medieval town | K49 | 13-14th c. | | | | | 249.0 | 241.2 | 47.1 | 27.2 |
| Kernavė medieval town | K50 | 13-14th c. | | | | | 228.0 | 220.0 | 41.3 | 24.4 |
| Kernavė medieval town | K51 | 13-14th c. | | | | | 229.0 | 221.0 | 41.0 | 24.5 |

**Table S5.** Sample sizes (*n*) of reference domestic horses (*Equus f. caballus/E. caballus*) and Przewalski's horses (*E. (f.) przewalskii*) with known heights, reconstructed heights, the total number of samples used for height estimation (known and reconstructed heights) and with known body masses. Note that numbers add up correctly only from left to right, as the height and body masses were obtained often from the same horse individuals.

| | **Domestic horses (*n*)** | **Przewalski's horses (*n*)** | **Combined sample (*n*)** |
|---|---|---|---|
| Individuals with known height | 15 | 0 | 15 |
| Individuals with reconstructed height | 42 | 46 | 88 |
| Individuals with either known or reconstructed height | 57 | 46 | 103 |
| Individuals with known body mass | 16 | 7 | 23 |
| Total number of individuals | 59 | 47 | 106 |

**Method S1**

Body size estimation

Most researchers still use the withers height – bone length ratios provided by May (1985: Tables 5-6) to estimate heights at the withers of horses and other equids. May computed these ratios by dividing the withers height and bone lengths of horses derived from a doctoral dissertation of Kiesewalter (1888), whose reference sample includes only 30 horses. Only one of these horses had a known living height. Kiesewalter reconstructed the heights of the 29 remaining horses from their skeletal dimensions (Kiesewalter 1888: Table 1). He thus used the so-called hybrid approach, which is increasingly used to estimate the stature of past humans represented only by skeletal dimensions and possessing likely somewhat different stature – limb bone length relationships than individuals in modern reference samples (see e.g., Ruff et al. 2012, Niskanen & Ruff 2018). This approach has been recently applied in estimating the heights of past horses and other *Equus* with updated reconstructions of heights from skeletal dimensions (Niskanen 2023; 2024; Rannamäe et al. 2025) and is also applied in this study, with reference samples including both domestic horses (*E. f. caballus*) and Przewalski's horses (*E. f. przewalskii*).

We estimated heights at the withers and at the croup, as well as body mass and the weight-carrying ability, following methods originally described in Niskanen (2023: Supplementary information B) and further refined in Niskanen (2024) in the case of height estimations. Changes in our reference sample size and composition allowed us to introduce revised equations to estimate heights and body masses. We also provide equations to estimate these measures of body size from phalangeal and distal tibial dimensions. Equations to estimate the so-called maximum rider weight from metapodial dimensions are from Niskanen (2023: Supplementary Information B). Names of skeletal dimensions used in this study are based on Eisenmann's (1986) naming system as applied in Niskanen (2023; 2014). Full names and abbreviations of these skeletal dimensions are provided in Table S1, which also provides abbreviations of these dimensions from Von Driesch (1976).

Reference samples presented in Table S5 used for generating equations to estimate heights and body masses was enlarged from those used in earlier studies (Niskanen 2023; 2024) with the addition of new specimens. These new specimens include skeletons of six Przewalski's horses (museum id: UN1354, UN1415, UN1539, UN1717, UN2015 and UN2020) and one mounted skeleton of a domestic horse measured at the Finnish Museum of Natural History in Helsinki. The final reference sample of 106 horses includes both domestic horses ($n$ = 59) and Przewalski's horses ($n$ = 47). To be included in this sample set, a horse had to have either living height, living body mass, or a complete enough skeleton (including an adequate number of phalangeal lengths) to reconstruct heights anatomically. Two horses with known living heights had missing phalangeal lengths and distal articular breadths of metapodials. One horse with body mass did not have living height and enough skeletal elements to reconstruct height, and two data points with body mass (domestic draft horses and one of seven data points for Przewalski's horses) are represented by mean values. Sample sizes range within 80–100 horses in height estimations and within 14–23 horses in body mass estimations.

Heights of 88 horses without known living heights but with complete enough skeletons were anatomically reconstructed as originally described in Niskanen (2023: Supplementary information B) and further refined in Niskanen (2024) based on 15 horses with both living dimensions and skeletal dimensions. A high correlation between anatomically reconstructed and actual withers heights ($r$ = 0.995, $n$ = 15) indicates that reconstructed heights are reasonably reliable proxies of living heights (Niskanen 2023: Supplementary information B).

As noted in Niskanen (2023, 2024), the joint breadth – bone length ratios of metacarpals and metatarsals reflect the relative lengths of these distal limb segments relative to the total height at the withers and at the croup. Similarly, relatively slender phalanges for their length tend to be relatively long, and relatively robust ones relatively short for these total heights. Therefore, the heights of horses were estimated with two-predictor equations from metapodial and phalangeal greatest lengths and greatest distal breadth-greatest length ratios. For this purpose, the following percentages of greatest distal breadths of greatest lengths were computed for metacarpals (MC11/MC1x100), metatarsals (MT11/MT1x100), first phalanges of forelimbs (P1fore6/P1fore1x100), first phalanges of hindlimbs (P1hind6/P1hind1x100), second phalanges of forelimbs (P2fore6/P2fore1x100), and second phalanges of hindlimbs (P2hind6/P2hind1x100). Regression equations to estimate withers height and croup height are provided in Table S6 and Table S7, respectively.

A reference sample used for generating regression equations to estimate body mass presented in Table S8 was based on that used in Niskanen (2023), but asses and zebras were excluded, and four Przewalski's horses (museum id: UN1354, UN1717, UN2015 and UN2020), whose bones were measured at the Finnish Museum of Natural History in Helsinki in 2024, were added. Due to the exclusion of asses and zebras, there is thus no need to correct estimates for horses downward between about two and five percent, depending on a predictor dimension in question, as was done in Niskanen (2023: Table S12). Larger sample sizes of equids with living body masses and skeletal measurements are needed to determine if the body mass – distal joint size relationships actually differ between different subgenera of *Equus* or if the difference noted in Niskanen (2023) was due to too small sample sizes of asses and zebras.

Proxies of distal metapodial joint size represented by products of distal articular breadths and least depths of the medial condyle were preferentially used in body mass estimations. MC11xMC13 is a proxy for the distal metacarpal joint size, and MT11xMT13 for the distal metatarsal joint size. Body mass of Lithuanian horses was estimated from these two variables. Estimates for four Lithuanian horses from both metacarpals and metatarsals were averaged. Body masses of wild horses from Ivanovskaya were estimated from MC11 (two horses), MT11 (two horses), P1fore4 (10 horses), P1hind4 (5 horses), P2fore4 (6 horses) or P2hind4 (6 horses) as predictor variables. Estimates from P2fore4 and P2hind4 were averaged because it was not known if these six second phalanges were from the forelimb or the hindlimb.

The maximum rider weight (XRW) refers to the total combined weight of the rider, the riding tack, and any other weight (including clothing, weapons, etc.) the horse should be able to carry in cavalry manoeuvres or trail riding if everything else (e.g., age, sex, condition, etc.) is held constant. These XRWs are expected to be about 10% heavier than recommended maximum weights for horses to carry, for example, in riding schools and trail riding. The method to estimate this XRW from metapodial dimensions is described in Niskanen (2023: Supplementary Information B), which also provides anatomical equations for these estimations. In this study, we apply the following two equations to archaeological horses from Lithuania:

$$\text{XRW (kg)} = [(\text{MC13} \times \text{MC11xMC13}^{0.84335}) / \text{MC2}] \times 1.60535$$

$$\text{XRW (kg)} = [(\text{MT13} \times \text{MT11xMT13}^{0.86094}) / \text{MT2}] \times 1.65952$$

The percent prediction error (%PE) values – computed as [(true-estimated) / estimated] × 100 – presented in Table S9 indicate the directional estimation error of estimates provided by regression equations. Known body masses and both known and anatomically reconstructed heights represent “true” values in these computations. Positive values indicate underestimation, and negative values indicate overestimation. Applying ANOVA on these %PE values indicated that there was some estimation bias related to subspecies in estimating heights (Table S9). Therefore, estimated heights are adjusted as in Niskanen (2024) by multiplying estimated heights by correction values presented in Table S10, derived by dividing “true” heights (known and reconstructed heights) by heights estimated with regression equations.

Equations to estimate heights and body mass were applied to 25 metapodial and phalangeal specimens dated to 5900–3800 BCE from the Neolithic site of Ivanovskaya in the easternmost parts of the Pontic-Caspian Steppe. These skeletal elements and their measurements are provided in Table S2. These equations were also applied to heights and body mass of Lithuanian horses predating the Common Era (CE) from Mineikiškės and the equation to assess body mass to a wild horse from Šventoji 43 and Late Bronze Age – Early Roman horses from Antilgė (Table S3). The same equations, as well as those for estimating the maximum rider weight, were also applied to metapodials of 43 Lithuanian horses divided into three temporal samples: pre-Viking period Iron Age ($n$ = 8), Viking period ($n$ = 9), and medieval period ($n$ = 26). These Lithuanian specimens and their metapodial measurements are provided in Table S4.

Height and body mass estimates for the Neolithic horses from Ivanovskaya are provided in Table S13. These horses are given height estimates adjusted for both domestic horses and Przewalski's horses because it is not known if these wild horses, which lived in the easternmost regions of the Pontic Caspian Steppe, were more similar to domestic horses or Przewalski's horses in the height – bone length relationship. These Neolithic horses from the easternmost parts of the Pontic-Caspian Steppe averaged somewhat taller and heavier than more recent horses from the western Pontic-Caspian Steppe represented by the Eneolithic horses of the Stredni Stog culture from Deriivka, as well as horses of the Eneolithic Tersek and Botai cultures of the northern Kazakh Steppe. Heights and body masses of these Eneolithic horses, which were included in Niskanen (2023), are re-estimated from their metacarpal dimensions by equations introduced in this study (Table S14).

As mean values in Table S14 show, equations introduced in this study give Eneolithic horses from Deriivka, Kozhai, Kumkeshu and Botai heights very similar withers heights and body masses than those used in Niskanen (2023), especially if we take into account the fact that the earlier estimates of withers heights were not adjusted for subspecies. This finding is expected because reference samples and methods used in this new study were developed from those used in Niskanen (2023). The Ivanovskaya horses were here estimated to have averaged 141.1–142.1 cm in height and 410.3 kg in weight not having noticeable lower heights at the withers than at the croup (see Table S13), which reflects the fact that heights of 19 out of 25 horses from this site are estimated from phalangeal dimensions. In any event, these wild horses, most likely belonging to a genetic cluster named NEO-NCAS (Neolithic North-Caspian) by Librado et al. (2021), were relatively large in size and thus provided a lot of meat for human societies hunting them. Wild horses were one of the most important game animals for hunter-gatherers and early herders, who lived between the lower Volga and Ural rivers (Vybornov et al., 2018: Table 1).

Height estimates of the Lithuanian horses are adjusted for domestic horses only. The adjusted height estimates, if applicable, and body masses of Lithuanian horses pre-dating Common Era (CE), are presented in Table S15. For later domestic Lithuanian horses, the adjusted height estimates, as well as estimated body masses and maximum rider weights, are provided separately for three temporal samples. Average heights of Lithuanian horses provided by equations introduced in this study are 127.4 cm for horses predating the Viking Age, 124.5 cm for those dated to the Viking Age, and 125.0 cm for those dated to the medieval period (Tables S16-S18). Averages of their body masses are 306.6 kg, 276.3 kg and 269.6 kg, respectively (Tables S16-S18). Their maximum rider weights (XRWs) estimated by applying equations from Niskanen (2023: Supplementary Information) are 92.1 kg, 85.8 kg and 79.7 kg, respectively (Tables S16-S18).

Estimating heights following May (1985) provides average withers heights of 127.6 cm, 125.4 cm and 127.1 cm, respectively, for these three temporal samples of Lithuanian horses (Tables S19-S21). Equations introduced in this study give horses with relatively broad joints for their metapodial lengths higher estimates than those of May (1985), whereas the opposite occurs for horses with narrow joints for their metapodial lengths. This finding is highly expected because, as noted in Niskanen (2023; 2024), the joint breadth – bone length ratios of metapodials reflect the relative lengths of these distal limb segments relative to the total height at the withers and at the croup.

In any event, the Lithuanian horses included in this study averaged shorter than medieval European horses from western and central Europe, which were generally 130-140 cm tall (Benkert 2023) as estimated following May (1985). The weight-carrying ability of these small horses of about the same size and perhaps the overall build as the current Gotland Russ ponies is thus of some importance because it has obvious implications for their usability for riding as well as for their use as packhorses. Their average maximum rider weights (XRWs) provided in tables 12a, b and c are quite expected for horses of their size, but there is quite a lot of variation in these values. Metapodials of some horses indicate that they could have carried a rider weighing considerably more than 100 kg. Therefore, some of these horses could have been suitable mounts for knights and other armoured men-at-arms based on their metapodial dimensions, but whether the backs of these horses were able to tolerate this much weight is unknown. The true weight-carrying ability of a horse is determined by its weakest point (Niskanen 2023: Supplementary information).

**Table S6**. Regression equations to estimate withers height in horses. Correlation coefficients (*r*), standard errors of estimates (SEE) and sample sizes (*n*) are provided. All dimensions are in millimeters. See Table S1 for the full names of the abbreviations of variables.

| **Equations to estimate withers height** | ***r*** | **SEE** | ***n*** |
|---|---|---|---|
| 6.121 × MC1 + 21.712 × MC11/MC1x100 – 492.999 | 0.974 | 43.063 | 100 |
| 5.526 × MT1 + 29.988 × MT11/MT1x100 – 668.228 | 0.978 | 39.803 | 100 |
| 15.411 × P1fore1 + 4.661 × P1fore6/P1fore1x100 – 197.933 | 0.985 | 31.446 | 88 |
| 14.942 × P1hind1 + 1.316 × P1hind6/P1hind1x100 + 74.560 | 0.978 | 38.478 | 86 |
| 28.727 × P2fore1 + 5.5.525 × P2fore6/P2fore1x100 – 586.212 | 0.969 | 46.129 | 84 |
| 26.716 × P2hind1 + 5.439 × P2hind6/P2hind1x100 – 461.135 | 0.962 | 52.012 | 80 |

**Table S7**. Regression equations to estimate croup height in horses. Correlation coefficients (*r*), standard errors of estimates (SEE) and sample sizes (*n*) are provided. All dimensions are in millimeters. See Table S1 for the full names of the abbreviations of variables.

| **Equations to estimate croup height** | ***r*** | **SEE** | ***n*** |
|---|---|---|---|
| 5.657 × MC1 + 20.236 × MC11/MC1x100 – 347.441 | 0.982 | 33.107 | 100 |
| 5.105 × MT1 + 27.540 × MT11/MT1x100 – 501.891 | 0.985 | 29.719 | 100 |
| 14.030 × P1fore1 + 3.695 × P1fore6/P1fore1x100 – 20.892 | 0.989 | 24.637 | 88 |
| 13.641 × P1hind1 + 0.864 × P1hind6/P1hind1x100 + 211.711 | 0.983 | 30.354 | 86 |
| 26.357 × P2fore1 + 5.007 × P2fore6/P2fore1x100 – 412.081 | 0.977 | 36.707 | 84 |
| 24.416 × P2hind1 + 5.073 × P2hind6/P2hind1x100 – 308.009 | 0.962 | 47.465 | 80 |

**Table S8.** Regression equations to estimate body mass in horses. Correlation coefficients (*r*), standard errors of estimates (SEE) and sample sizes (*n*) are provided. Predictor values are in millimeters. Body mass in kilograms. See Table S1 for the full names of the abbreviations of variables.

| **Equation to estimate body mass** | ***r*** | **SEE** | ***n*** |
|---|---|---|---|
| 0.309 × MC11xMC13 – 97.592 | 0.975 | 43.153 | 23 |
| 0.312 × MT11xMT13 – 91.884 | 0.973 | 44.872 | 23 |
| 18.611 × MC11 – 554.059 | 0.967 | 48.960 | 23 |
| 18.958 × MT11 – 562.329 | 0.967 | 49.644 | 23 |
| 17.902 × P1fore4 – 605.852 | 0.957 | 47.210 | 14 |
| 17.033 × P1hind4 – 567.697 | 0.947 | 52.488 | 14 |
| 19.765 × P2fore4 – 669.299 | 0.946 | 52.539 | 14 |
| 19.481 × P2hind4 – 653.411 | 0.944 | 53.552 | 14 |
| 0.147 × TIB7xTIB8 – 154.973 | 0.994 | 18.856 | 7 |

**Table S9.** Percent prediction error (%PE) values with mean and sample size (*n*) provided, as well as ANOVA *F* computed on these %PE values and significance (sig.). See Table S1 for the full names of the variable abbreviations.

| | ***caballus*** | | ***przewalskii*** | | **ANOVA** | |
|---|---|---|---|---|---|---|
| Percent prediction errors | Mean | *n* | Mean | *n* | *F* | Sig. |
| WH from metacarpal | 1.3129 | 54 | -1.5549 | 46 | 21.979 | <0.001 |
| WH from metatarsal | 1.0576 | 54 | -1.2365 | 46 | 15.785 | <0.001 |
| WH from P1fore | 0.0679 | 45 | -0.1111 | 43 | 0.102 | 0.751 |
| WH from P1hind | -0.4573 | 44 | 0.4171 | 42 | 1.682 | 0.198 |
| WH from P2fore | -0.3877 | 44 | 0.3441 | 40 | 0.759 | 0.386 |
| WH from P2hind | -1.1431 | 41 | 1.1148 | 39 | 5.730 | 0.019 |
| CH from metacarpal | 0.8402 | 54 | -0.9988 | 46 | 14.594 | <0.001 |
| CH from metatarsal | 0.6239 | 54 | -0.7295 | 46 | 9.437 | 0.003 |
| CH from P1fore | -0.3467 | 45 | 0.3290 | 43 | 2.468 | 0.120 |
| CH from P1hind | -0.8100 | 44 | 0.7994 | 42 | 10.151 | 0.002 |
| CH from P2fore | -0.7398 | 44 | 0.7446 | 40 | 5.340 | 0.023 |
| CH from P2hind | -1.4639 | 41 | 1.4685 | 39 | 13.367 | <0.001 |
| BM from MC11xMC13 | -1.0016 | 14 | 0.4682 | 7 | 0.067 | 0.799 |
| BM from MT11xMT13 | -4.3105 | 14 | 6.6936 | 7 | 4.070 | 0.058 |
| BM from MC11 | 2.7538 | 14 | -1.4066 | 7 | 0.198 | 0.661 |
| BM from MT11 | -0.9020 | 14 | 3.5545 | 7 | 0.876 | 0.362 |
| BM from P1fore4 | 5.8182 | 10 | 0.8490 | 4 | 0.084 | 0.778 |
| BM from P1hind4 | 0.3242 | 10 | 8.0318 | 4 | 0.324 | 0.580 |
| BM from P2fore4 | 11.9534 | 10 | -5.4050 | 4 | 0.589 | 0.457 |
| BM from P2hind4 | 12.6766 | 10 | -5.2862 | 4 | 0.551 | 0.472 |

**Table S10.** Mean ratios between observed or reconstructed heights and estimated heights. Values greater than 1 indicate underestimation, and those smaller than 1 indicate overestimation. ANOVA *F* and significance (sig.) are also provided. See Table S1 for the full names of the abbreviations of variables.

| | **Species/subspecies** | | **ANOVA** | |
|---|---|---|---|---|
| | *caballus* | *prezewalskii* | *F* | Sig. |
| WH from MC | 1.0134 | 0.9841 | 21.979 | <0.001 |
| WH from MT | 1.0107 | 0.9872 | 15.785 | <0.001 |
| WH from P1fore | 1.0014 | 0.9984 | 0.102 | 0.751 |
| WH from P1hind | 0.9970 | 1.0033 | 1.682 | 0.198 |
| WH from P2fore | 0.9980 | 1.0023 | 0.759 | 0.386 |
| WH from P2hind | 0.9908 | 1.0102 | 5.730 | 0.019 |
| CH from MC | 1.0086 | 0.9899 | 14.594 | <0.001 |
| CH from MT | 1.0063 | 0.9925 | 9.437 | 0.003 |
| CH from P1fore | 0.9972 | 1.0030 | 2.468 | 0.120 |
| CH from P1hind | 0.9931 | 1.0075 | 10.151 | 0.002 |
| CH from P2fore | 0.9940 | 1.0068 | 5.340 | 0.023 |
| CH from P2hind | 0.9873 | 1.0140 | 13.367 | <0.001 |

**Data S1**

Description of the archaeological context of Levänluhta, Finland

Levänluhta in Isokyrö is an Iron Age water burial site containing uncremated human remains dated to c. 300–800 CE. The excavations, from the 1800s until the 1980s, have revealed commingled human remains from 98 individuals, buried alongside artefacts and animal bones (Wessman, 2009). The Levänluhta water burial is unique as the norm of Iron Age Finland was to cremate the dead, but here, 98 humans, mainly females (Maijanen et al., 2021), were buried in a small lake with artefacts (Wessman, 2009). The artefact finds include precious copper alloy brooches, arm rings and other dress implements, as well as a Provincial Roman copper cauldron (Wessman, 2009). Animal bones have also been excavated, with the most common species including horse and cattle, with single finds of sheep, dog, chicken and probably a wild goose; however, most of the animal finds, including the horses, date to the Finnish Medieval or Post-Medieval period (1300–1800 CE) (Wessman, 2018). Thus, most of the animal finds were not part of the burial practices but rather reflect later agrarian use of the site (Wessman et al., 2018). Wessman et al. (2018) argue that it is possible that the people burying the domestic animals may have been aware of the site's sacred history. Samples analysed for this study were excavated in 1912 by A.M Tallgren (KM6110:7.701; maxillary tooth) and in 1982 by Aarni Erä-Esko (KM21814; mandibular left M3, mandibular right P2, right tibia and right radius). Our analysed horse bones have been dated to the 16th century CE (KM5111:7.701), and 14th century CE (KM21814:23,525, 526 and 584).

**Table S11.** Overlapping primer pairs used to amplify a 572-573 bp base pair fragment of the mitochondrial control region from ancient subfossil horses (*Equus caballus*). Primers are from Cieslak et al. (2010), with the reverse primer reverse-complemented to present it in 5'-3' orientation.

| Primer | Primer sequence 5'-3' | Annealing temperature (°C) | Product size (bp) |
|---|---|---|---|
| HP1F | CTTCCCCTAAACGACAACAA | 47 | 116 bp |
| HP1R | CGAYGTACATAGGCCATTC | | |
| HP2F | CCCCCAYATAACACYATACC | 47 | 153 bp |
| HP2R | GGGGTATGCACGATYAATA | | |
| HP3F | GCCCCATGAATAATAAGCA | 47 | 178 bp |
| HP3R | CACGTAGTTGRGAGGGTTG | | |
| HP4F | ATCACARCCCATGTTCCAC | 47 | 153 bp |
| HP4R | ATGGCCCTGAAGAAAGAACC | | |
| HP5F | AAACGTGGGGGTTTCTAC | 47 | 136 bp |
| HP5R | GTGAGCATGGGCTGATTA | | |
| HP6F | GCCCATTCTTTCCCCTTA | 47 | 137 bp |
| HP6R | CTTTGACGGCCATAGCTG | | |
| HP7F | ATTTGGTATYTTTTATATTTGG | 47 | 116 bp |
| HP7R | GCTGATGCGGAGGRATA | | |

**Method S2**

PCR

We amplified and sequenced a 573 bp part of the mitochondrial control region using an overlapping primer set (primer set 2, HP1-HP7) from Cieslak et al. (2010), except that the reverse primers were reverse-complemented (Table S11), as the original publication erroneously reported them in 3' 5' orientation. First, we used the primer pair HP1F and HP1R to screen the samples for amplifiable mtDNA, and proceeded with the remaining primer pairs only if amplification was observed on a 2% agarose gel with the HP1F/R primer pair. PCR reactions were performed in 12.5 µl reaction volumes under the following conditions: 1 x PCR buffer (HotStarTaq, Qiagen), 0.2 mM of each dNTPs, 0.2 µM of F- and R-primers, 2.5 mM $MgCl_2$, 1 mg/mL BSA (Bovine serum albumin), 4 U/reactions of HotStarTaq DNA Polymerase (Qiagen), and 1 µl of template DNA. The thermal profile consisted of an initial denaturation at 95 ºC for 15 min, followed by 55 cycles of 94 ºC for 30 s, 47 ºC for 30 s and 72 ºC for 30 s, with a final extension at 72 ºC for 10 min. Each PCR included a negative control containing water instead of template DNA.

Sequence analysis

Furthermore, we compared the ancient horse sequences with those of modern breeds. We used an extensive dataset of Finnhorses (*n* = 743; Finnish native breed) (GenBank accession numbers: MN070243-MN070985) and other breeds (*n* = 118; MN070986-MN071106) (Kvist et al., 2019), as well as native breeds of eastern origin including Akhal-Teke (*n* = 18), Mesenskaya (*n* = 18), Mongolian (*n* = 16), Orlov (*n* = 18), Vyatskaya (*n* = 18) and Yakut (*n* = 20) (DQ327950-DQ328057; McGahern et al., 2006). In addition, we included sequences from Achilli et al. (2012), which were also used in the previous network analysis. All sequences were trimmed to 556 bp from the 5' end, because some modern sequences were slightly shorter. The dataset included only two Estonian Native Horse sequences (MN070986 and MN070987), and no Žemaitukas (Lithuanian native horse) sequences were available for comparison.

**Table S12.** Ancient wild and domestic horse (*Equus caballus/E. ferus*) sequences used for comparison. GenBank accession numbers, sample name, site and dating (based on $^{14}$C dating when available, otherwise based on archaeological context), haplogroup determined in this study following the nomenclature of Achilli et al. (2012), and literature references are provided. A question mark (?) indicates samples for which the haplogroup could not be determined.

| Species name | GenBank accession number | Sample | Site | Dating | Haplogroup | Reference | DOI |
|---|---|---|---|---|---|---|---|
| *Equus caballus* | OR067839 | XSS01H | Shihuyao tombs, Yilan Habir Mountain, China | 389–208 cal BCE | B | Zhu et al. 2024 | https://doi.org/10.3390/genes15060790 |
| *Equus ferus* | FJ204324 | SP1181D | Bol'shoy Lyakhovsky Isl., North Siberia, Russia | Pleistocene | F | Cieslak et al. 2010 | https://doi.org/10.1371/journal.pone.0015311 |
| *Equus ferus* | FJ204358 | Orl4 | Orlovka, Moldova | 4000 BCE, Eneolithic | N | Cieslak et al. 2010 | https://doi.org/10.1371/journal.pone.0015311 |
| *Equus ferus* | FJ204359 | May1 | Mayaki, Ukraine | 3600-3100 BCE, Copper Age | N | Cieslak et al. 2010 | https://doi.org/10.1371/journal.pone.0015311 |
| *Equus ferus* | FJ204364 | May4 | Mayaki, Ukraine | 3600-3100 BCE, Copper Age | N | Cieslak et al. 2010 | https://doi.org/10.1371/journal.pone.0015311 |
| *Equus ferus* | FJ204384 | 44 | Atxoste, Spain | 5500-4950 BCE, Mesolithic | ? | Cieslak et al. 2010 | https://doi.org/10.1371/journal.pone.0015311 |

| | | | | | | | |
|---|---|---|---|---|---|---|---|
| *Equus ferus* | HM802276 | 2 | Cueva Fosca - Valencia-Cartellon, Spain | 5200-4900 BCE, Copper Age and Mesolithic | ? | Cieslak et al. 2010 | https://doi.org/10.1371/journal.pone.0015311 |
| *Equus caballus* | HM802277 | 20 | Cueva Rubia-Valmayor/Madrid, Spain | 2880-2570 BCE, Copper Age and Mesolithic | ? | Cieslak et al. 2010 | https://doi.org/10.1371/journal.pone.0015311 |
| *Equus caballus* | HM802278 | 21 | Cueva Rubia-Valmayor/Madrid, Spain | 2900-2500 BCE, Copper Age and Mesolithic | ? | Cieslak et al. 2010 | https://doi.org/10.1371/journal.pone.0015311 |
| *Equus ferus* | HM802280 | 35 | Cueva Fosca - Valencia-Cartellon, Spain | 5380-5210 BCE, Copper Age and Mesolithic | ? | Cieslak et al. 2010 | https://doi.org/10.1371/journal.pone.0015311 |
| *Equus caballus* | HM802281 | 27 | El Caprichio-Madrid, Spain | 4300-2200 BCE, Copper Age and Mesolithic | ? | Cieslak et al. 2010 | https://doi.org/10.1371/journal.pone.0015311 |
| *Equus caballus* | DQ900924 | K316 | Dashanqian site, Inner Mongolia, China | 2800 BP | P | Cai et al. 2007 | https://doi.org/10.1080/10020070708541034 |
| *Equus caballus* | DQ900925 | K420 | Dashanqian site, Inner Mongolia, China | 2800 BP | F | Cai et al. 2007 | https://doi.org/10.1080/10020070708541034 |

| | | | | | | | |
|---|---|---|---|---|---|---|---|
| *Equus caballus* | AY049720 | | Kwakji archaeological site, Jeju, Korea | 700-800 CE | ? | Jung et al. 2002 | https://doi.org/10.1016/S1016-8478(23)15096-5 |
| *Equus sp.* | DQ007556 | JW175.1 | Hohlefels, Germany | 12550±60 BP | ? | Weinstock et al. 2005 | https://doi.org/10.1371/journal.pbio.0030241 |
| *Equus caballus* | OM222617 | SNU-A001 | Gongpyeongdong site, Korea | 15th century CE | Q | Hong et al. 2022 | https://doi.org/10.5713/ab.21.0500 |
| *Equus caballus* | MW534079 | IMCBSB RAS Mon Gan1 | Bulgan aimag, valley of the river, Mongolia | late 12th to mid-10th century BCE | M | Kusliy et al. 2021 | https://doi.org/10.3390/genes12030412 |
| *Equus caballus* | MW534080 | IMCBSB RAS Mon Gan3 | Bulgan aimag, valley of the river, Mongolia | late 12th to mid-10th century BCE | M | Kusliy et al. 2021 | https://doi.org/10.3390/genes12030412 |
| *Equus caballus* | MW534081 | IMCBSB RAS Mon Gan11 | Bulgan aimag, valley of the river, Mongolia | late 12th to mid-10th century BCE | G | Kusliy et al. 2021 | https://doi.org/10.3390/genes12030412 |
| *Equus caballus* | MW534082 | IMCBSB RAS Mon Gan14 | Bulgan aimag, valley of the river, Mongolia | late 12th to mid-10th century BCE | L | Kusliy et al. 2021 | https://doi.org/10.3390/genes12030412 |
| *Equus caballus* | MW534083 | IMCBSB RAS Mon Gan18 | Bulgan aimag, valley of the river, Mongolia | late 12th to mid-10th century BCE | L | Kusliy et al. 2021 | https://doi.org/10.3390/genes12030412 |

| | | | | | | | |
|---|---|---|---|---|---|---|---|
| *Equus caballus* | PP700507 | EQ005 | Żółte, Western Pomerania, Lower Oder, Poland | 995 - 1158 cal CE | M | Popovic et al. 2024 | https://doi.org/10.1016/j.jasrep.2024.104530 |
| *Equus caballus* | PP700508 | EQ010 | Ostrów Lednicki 2, Greater Poland, Poland | 1022 - 1159 cal CE | L | Popovic et al. 2024 | https://doi.org/10.1016/j.jasrep.2024.104530 |
| *Equus caballus* | PP700509 | EQ016 | Chycina 19, Lubusz Land, Lower Oder, Poland | 1027 - 1166 cal CE | B | Popovic et al. 2024 | https://doi.org/10.1016/j.jasrep.2024.104530 |
| *Equus caballus* | PP700510 | EQ019 | Kałdus 3, Chełmno Land, Lower Vistula, Poland | 992 - 1154 cal CE | B | Popovic et al. 2024 | https://doi.org/10.1016/j.jasrep.2024.104530 |
| *Equus caballus* | PP700511 | EQ020 | Kałdus 3, Chełmno Land, Lower Vistula, Poland | early 12th | E | Popovic et al. 2024 | https://doi.org/10.1016/j.jasrep.2024.104530 |
| *Equus caballus* | PP700512 | EQ024 | Kałdus 3, Chełmno Land, Lower Vistula, Poland | 11th/12th | G | Popovic et al. 2024 | https://doi.org/10.1016/j.jasrep.2024.104530 |
| *Equus caballus* | PP700513 | EQ025 | Kałdus 3, Chełmno Land, Lower Vistula, Poland | 11th/12th | M | Popovic et al. 2024 | https://doi.org/10.1016/j.jasrep.2024.104530 |

| | | | | | | | |
|---|---|---|---|---|---|---|---|
| *Equus caballus* | PP700514 | EQ026 | Kałdus 3, Chełmno Land, Lower Vistula, Poland | 1039 - 1210 cal CE | M | Popovic et al. 2024 | https://doi.org/10.1016/j.jasrep.2024.104530 |
| *Equus caballus* | PP700515 | EQ027 | Gdańsk 1, Grodzka, eastern Pomerania, Lower Vistula, Poland | 1030 - 1198 cal CE | M | Popovic et al. 2024 | https://doi.org/10.1016/j.jasrep.2024.104530 |
| *Equus caballus* | PP700516 | EQ029 | Gdańsk 1, Grodzka, eastern Pomerania, Lower Vistula, Poland | 1042 - 1219 cal CE | M | Popovic et al. 2024 | https://doi.org/10.1016/j.jasrep.2024.104530 |
| *Equus caballus* | PP700517 | EQ032 | Szczecin Podzamcze, Western Pomeria, Lower Oder, Poland | 13th | B | Popovic et al. 2024 | https://doi.org/10.1016/j.jasrep.2024.104530 |
| *Equus caballus* | PP700518 | EQ033 | Bydgoszcz 1, Greater Poland, Poland | late 11th | K | Popovic et al. 2024 | https://doi.org/10.1016/j.jasrep.2024.104530 |
| *Equus caballus* | PP700519 | EQ036 | Ostrów Lednicki 2 (Rybitwy), Greater Poland, Poland | 9th - 13th/14th | L | Popovic et al. 2024 | https://doi.org/10.1016/j.jasrep.2024.104530 |
| *Equus caballus* | PP700520 | EQ046 | Kałdus 3, Chełmno Land, Lower Vistula, Poland | 992 - 1154 cal CE | L | Popovic et al. 2024 | https://doi.org/10.1016/j.jasrep.2024.104530 |

| | | | | | | | |
|---|---|---|---|---|---|---|---|
| *Equus caballus* | PP700521 | EQ061 | Kałdus 3, Chełmno Land, Lower Vistula, Poland | 1035 - 1210 cal CE | E | Popovic et al. 2024 | https://doi.org/10.1016/j.jasrep.2024.104530 |
| *Equus caballus* | PP700522 | EQ062 | Starorypin 3, Chełmno Land, Lower Vistula, Poland | 1030 - 1198 cal CE | L | Popovic et al. 2024 | https://doi.org/10.1016/j.jasrep.2024.104530 |
| *Equus caballus* | PP700523 | EQ067 | Kruszwica 4, Kuyavia, Greater Poland, Poland | early 12th | I | Popovic et al. 2024 | https://doi.org/10.1016/j.jasrep.2024.104530 |
| *Equus caballus* | PP700524 | EQ068 | Kruszwica 4 m. 365, Kuyavia, Greater Poland, Poland | 9th - early 14th | R | Popovic et al. 2024 | https://doi.org/10.1016/j.jasrep.2024.104530 |
| *Equus caballus* | PP700525 | EQ069 | Kruszwica 4, Kuyavia, Greater Poland, Poland | 1042 - 1219 cal CE | G | Popovic et al. 2024 | https://doi.org/10.1016/j.jasrep.2024.104530 |
| *Equus caballus* | PP700526 | EQ070 | Kruszwica 4, Kuyavia, Greater Poland, Poland | 1165 - 1267 cal CE | B | Popovic et al. 2024 | https://doi.org/10.1016/j.jasrep.2024.104530 |
| *Equus caballus* | PP700527 | EQ071 | Kruszwica 4, Kuyavia, Greater Poland, Poland | 9th - early 14th | G | Popovic et al. 2024 | https://doi.org/10.1016/j.jasrep.2024.104530 |
| *Equus caballus* | PP700528 | EQ072 | Kruszwica 4, Kuyavia, Greater Poland, Poland | 9th - early 14th | G | Popovic et al. 2024 | https://doi.org/10.1016/j.jasrep.2024.104530 |

| | | | | | | | |
|---|---|---|---|---|---|---|---|
| *Equus caballus* | PP700529 | EQ073 | Kruszwica 4, Kuyavia, Greater Poland, Poland | 1033 - 1206 cal CE | P | Popovic et al. 2024 | https://doi.org/10.1016/j.jasrep.2024.104530 |
| *Equus caballus* | PP700530 | EQ074 | Kruszwica 4, Kuyavia, Greater Poland, Poland | late 10th - 13th | I | Popovic et al. 2024 | https://doi.org/10.1016/j.jasrep.2024.104530 |
| *Equus caballus* | PP700531 | EQ075 | Kruszwica 4, Kuyavia, Greater Poland, Poland | 9th - early 14th | L | Popovic et al. 2024 | https://doi.org/10.1016/j.jasrep.2024.104530 |
| *Equus caballus* | PP700534 | EQ080 | Ostrów Lednicki 2, Greater Poland, Poland | 899 - 1153 cal CE | M | Popovic et al. 2024 | https://doi.org/10.1016/j.jasrep.2024.104530 |
| *Equus caballus* | PP700537 | EQ091 | Górzyca 20, Lubusz Land, Poland | 892 - 1023 cal CE | D | Popovic et al. 2024 | https://doi.org/10.1016/j.jasrep.2024.104530 |
| *Equus caballus* | PP700538 | EQ092 | Gdańsk, st. 1 (Sukiennicza), Eastern Pomerania, Lower Vistula Poland | 1025 - 1160 cal CE | F | Popovic et al. 2024 | https://doi.org/10.1016/j.jasrep.2024.104530 |
| *Equus caballus* | PP700542 | EQ098 | Kołobrzeg - Budzistowo 1, Western Pomerinia, Lower Oder, Poland | 1045 - 1225 cal CE | L | Popovic et al. 2024 | https://doi.org/10.1016/j.jasrep.2024.104530 |

| | | | | | | | |
|---|---|---|---|---|---|---|---|
| *Equus caballus* | PP700543 | EQ099 | Kołobrzeg - Budzistowo 1, Western Pomerinia, Lower Oder, Poland | 1045 - 1225 cal CE | B | Popovic et al. 2024 | https://doi.org/10.1016/j.jasrep.2024.104530 |
| *Equus caballus* | PP700544 | EQ108 | Gdańsk, st. 1, Eastern Pomerania, Lower Vistula | 1045 - 1225 cal CE | M | Popovic et al. 2024 | https://doi.org/10.1016/j.jasrep.2024.104530 |
| *Equus caballus* | PP700545 | EQ109 | Gdańsk, st. 1, Eastern Pomerania, Lower Vistula | early 12th | L | Popovic et al. 2024 | https://doi.org/10.1016/j.jasrep.2024.104530 |
| *Equus caballus* | PP700546 | EQ110 | Gdańsk, st. 1, Eastern Pomerania, Lower Vistula | 1028 - 1172 cal CE | L | Popovic et al. 2024 | https://doi.org/10.1016/j.jasrep.2024.104530 |
| *Equus caballus* | PP700547 | EQ111 | Gdańsk, st. 1, Eastern Pomerania, Lower Vistula | 1033 - 1206 cal CE | L | Popovic et al. 2024 | https://doi.org/10.1016/j.jasrep.2024.104530 |
| *Equus caballus* | PP700549 | EQ113 | Gdańsk, st. 2, Eastern Pomerania, Lower Vistula | late 12th - early 13th | N | Popovic et al. 2024 | https://doi.org/10.1016/j.jasrep.2024.104530 |
| *Equus caballus* | PP700550 | EQ114 | Wolin - Miasto, st. 1, Western Pomerania, Lower Oder, Poland | 1028 - 1172 cal CE | B | Popovic et al. 2024 | https://doi.org/10.1016/j.jasrep.2024.104530 |

| | | | | | | | |
|---|---|---|---|---|---|---|---|
| *Equus caballus* | PP700552 | EQ116 | Wolin - Miasto, st. 1, Western Pomerania, Lower Oder, Poland | 1167 - 1269 cal CE | M | Popovic et al. 2024 | https://doi.org/10.1016/j.jasrep.2024.104530 |
| *Equus caballus* | PP700553 | EQ117 | Wolin - Miasto, st. 1, Western Pomerania, Lower Oder, Poland | 1030 - 1198 cal CE | E | Popovic et al. 2024 | https://doi.org/10.1016/j.jasrep.2024.104530 |
| *Equus caballus* | PP700554 | EQ120 | Ostrów Lednicki 2 (Rybitwy 3a/4), Greater Poland, Poland | 987 - 1153 cal CE | B | Popovic et al. 2024 | https://doi.org/10.1016/j.jasrep.2024.104530 |
| *Equus caballus* | PP700557 | EQ123 | Ostrów Lednicki 2 (1991), Greater Poland, Poland | 899 - 1147 cal CE | C | Popovic et al. 2024 | https://doi.org/10.1016/j.jasrep.2024.104530 |
| *Equus caballus* | PP700558 | EQ124 | Ostrów Lednicki 2, Greater Poland, Poland | 890 - 1020 cal CE | N | Popovic et al. 2024 | https://doi.org/10.1016/j.jasrep.2024.104530 |
| *Equus caballus* | PP700559 | EQ126 | Kruszwica 4, Kuyavia, Greater Poland, Poland | 9th - early 14th | ? | Popovic et al. 2024 | https://doi.org/10.1016/j.jasrep.2024.104530 |

| | | | | | | | |
|---|---|---|---|---|---|---|---|
| *Equus caballus* | PP700560 | EQ129 | Wolin - Miasto, st. 1/wykop 6, Western Pomerania, Lower Oder, Poland | early 10th | L | Popovic et al. 2024 | https://doi.org/10.1016/j.jasrep.2024.104530 |
| *Equus caballus* | PP700562 | EQ148 | Wolin, st. 3, Western Pomerania, Lower Oder, Poland | 1028 - 1162 cal CE | L | Popovic et al. 2024 | https://doi.org/10.1016/j.jasrep.2024.104530 |
| *Equus caballus* | PP700565 | EQ152 | Wolin, st. 3, Western Pomerania, Lower Oder, Poland | 1025 - 1160 cal CE | B | Popovic et al. 2024 | https://doi.org/10.1016/j.jasrep.2024.104530 |
| *Equus caballus* | PP700566 | EQ158 | Biskupin 15a, Greater Poland, Poland | 1490 - 1649 cal CE | L | Popovic et al. 2024 | https://doi.org/10.1016/j.jasrep.2024.104530 |
| *Equus caballus* | KT985979 | HUk1 | Altai Republic, plateau Ukok, funerary complex Ak-Alakha, Russia | 4th-3rd century BCE | I | Vorobieva et al. 2010 | https://doi.org/10.1371/journal.pone.0241997 |
| *Equus caballus* | KT985980 | HUk2 | Altai Republic, plateau Ukok, funerary complex Verkh-Kaldzhin-2, Russia | 4th-3rd century BCE | N | Vorobieva et al. 2010 | https://doi.org/10.1371/journal.pone.0241997 |

| | | | | | | | |
|---|---|---|---|---|---|---|---|
| *Equus caballus* | MK449357 | HUk3 | Altai mountains, Ukok plateau, kurgan group Ak-Alaha-II, Russia | 7th century BCE | Q | Vorobieva et al. 2010 | https://doi.org/10.1371/journal.pone.0241997 |
| *Equus caballus* | MK449358 | HUk4 | Altai mountains, Ukok plateau, kurgan group Ak-Alaha-II, Russia | 7th century BCE | A | Vorobieva et al. 2010 | https://doi.org/10.1371/journal.pone.0241997 |
| *Equus caballus* | MK467454 | HUk6 | Altai mountains, Ukok plateau, kurgan group Ak-Alaha-II, Russia | 7th century BCE | P | Vorobieva et al. 2010 | https://doi.org/10.1371/journal.pone.0241997 |
| *Equus ferus* | MK467455 | HD2 | Altai mountains, archaeological site Denisova cave, Russia | 30-50 kya | ? | Vorobieva et al. 2010 | https://doi.org/10.1371/journal.pone.0241997 |

**Table S13.** Estimated withers and croup heights and body masses of horses from Ivanovskaya, adjusted to *E. caballus* or *E. przewalskii.* Heights provided by regression equations in millimeters have been converted to centimeters. Body mass is given in kilograms.

| **Specimen** | **Bone** | **Withers height (*caballus*)** | **Croup height (*caballus*)** | **Withers height (*przewalskii*)** | **Croup height (*przewalskii*)** | **Body mass** |
|---|---|---|---|---|---|---|
| 1 | Metacarpal III | 137.9 | 138.2 | 133.9 | 135.6 | 357.9 |
| 2 | Metacarpal III | 134.7 | 135.2 | 130.8 | 132.7 | 357.9 |
| 3 | Metatarsal III | 146.5 | 146.0 | 143.1 | 143.2 | 499.3 |
| 4 | Metatarsal III | 139.5 | 139.6 | 136.2 | 136.9 | 442.4 |
| 5 | Phalanx I (fore) | 142.0 | 141.6 | 141.5 | 142.4 | 432.4 |
| 6 | Phalanx I (fore) | 143.7 | 143.1 | 143.3 | 144.0 | 396.6 |
| 7 | Phalanx I (fore) | 147.3 | 146.3 | 146.8 | 147.1 | 486.1 |
| 8 | Phalanx I (fore) | 149.8 | 148.6 | 149.3 | 149.5 | 486.1 |
| 9 | Phalanx I (fore) | 141.8 | 141.2 | 141.4 | 142.1 | 396.6 |
| 10 | Phalanx I (fore) | 134.7 | 134.9 | 134.3 | 135.7 | 360.8 |
| 11 | Phalanx I (fore) | 130.5 | 131.3 | 130.1 | 132.0 | 378.7 |
| 12 | Phalanx I (fore) | 152.2 | 151.0 | 151.7 | 151.9 | 450.3 |
| 13 | Phalanx I (fore) | 147.0 | 146.3 | 146.6 | 147.1 | 432.4 |
| 14 | Phalanx I (fore) | 132.3 | 132.9 | 131.9 | 133.6 | 325.0 |
| 15 | Phalanx I (hind) | 151.3 | 150.1 | 152.3 | 152.3 | 369.1 |
| 16 | Phalanx I (hind) | 138.8 | 138.5 | 139.7 | 140.5 | 437.3 |
| 17 | Phalanx I (hind) | 128.6 | 129.2 | 129.4 | 131.1 | 352.1 |
| 18 | Phalanx I (hind) | 138.7 | 138.4 | 139.6 | 140.4 | 437.3 |
| 19 | Phalanx I (hind) | 146.5 | 145.8 | 147.4 | 148.0 | 352.1 |

| | | | | | | |
|---|---|---|---|---|---|---|
| 20 | Phalanx II | 134.3 | 134.6 | 135.9 | 137.3 | 359.0 |
| 21 | Phalanx II | 142.0 | 141.6 | 143.7 | 144.4 | 417.9 |
| 22 | Phalanx II | 139.8 | 139.6 | 141.5 | 142.4 | 417.9 |
| 23 | Phalanx II | 146.5 | 145.7 | 148.2 | 148.6 | 476.8 |
| 24 | Phalanx II | 148.3 | 147.3 | 150.0 | 150.2 | 437.5 |
| 25 | Phalanx II | 141.6 | 141.2 | 143.3 | 144.0 | 398.3 |
| | n | 25 | 25 | 25 | 25 | 25 |
| | Mean | 141.5 | 141.1 | 141.3 | 142.1 | 410.3 |
| | Std. | 6.5 | 6.0 | 6.9 | 6.4 | 49.0 |
| | Min. | 128.6 | 129.2 | 129.4 | 131.1 | 325.0 |
| | Max. | 152.2 | 151.0 | 152.3 | 152.3 | 499.3 |

**Table S14.** Mean estimated heights (cm) and body masses (kg) of Neolithic horses from Ivanovskaya compared with Eneolithic period horses from the Eurasian steppe. Body sizes of the Eneolithic horses, which were included in Niskanen (2023: Table 1), are re-estimated with equations introduced in this current study to make them directly comparable.

| Sample | *n* | Withers height (*caballus*) | Croup height (*caballus*) | Withers height (*przewalskii*) | Croup height (*przewalskii*) | Body mass | Withers height (Niskanen 2023) | Body mass (Niskanen 2023) |
|---|---|---|---|---|---|---|---|---|
| Ivanovskaya | 25 | 141.5 | 141.1 | 141.3 | 142.1 | 410.3 | -- | -- |
| Deriivka (Stredni Stog) | 15 | 139.3 | 139.4 | 135.3 | 136.9 | 384.4 | 136.2 | 373.7 |
| Kozhai (Tersek culture) | 12 | 139.7 | 139.8 | 135.7 | 137.2 | 376.5 | 137.2 | 369.4 |
| Kumkeshu (Tersek culture) | 41 | 140.4 | 140.4 | 136.3 | 137.8 | 371.8 | 138.1 | 368.8 |
| Botai (Botai culture) | 18 | 140.7 | 140.7 | 136.6 | 138.1 | 364.4 | 138.9 | 362.8 |

**Table S15.** Estimated withers and croup heights and body masses of Lithuanian horses predating the Common Era (CE). Heights provided by regression equations in millimeters have been converted to centimeters. Body mass is given in kilograms.

| Site | Specimen | Date | Withers height (*caballus*) | Croup height (*caballus*) | Body mass |
|---|---|---|---|---|---|
| Šventoji 43 | aEca15* (Tibia) | 3881 BC ± 57 | -- | -- | 265.5 |
| Mineikiškės | Phalanx I (fore)** | Late Bronze Age | 117.9 | 119.5 | 300.0 |
| Mineikiškės | Phalanx I (hind)** | Late Bronze Age | 120.7 | 122.0 | 251.6 |
| Antilgė | Phalanx I (fore or hind?) | Late Bronze Age – Early Roman | -- | -- | 310.2 / 303.9 |
| Antilgė | Tibia | Late Bronze Age – Early Roman | -- | -- | 200.4 |

* See Table 1 in main text

** possibly the same individual

**Table S16.** Estimated withers and croup heights, body masses, and maximum rider weights (XRW) of Lithuanian horses predating the Viking Age. Heights provided by regression equations in millimeters have been converted to centimeters. Body mass and rider weight are given in kilograms.

| Site | Specimen | Date | Withers height (*caballus*) | Croup height (*caballus*) | Body mass | XRW |
|---|---|---|---|---|---|---|
| Pagrybis cemetary | 142 | 6-7th c. CE | 130.3 | 131.2 | 323.5 | 90.3 |
| Pagrybis cemetary | 145 | 601–662 CE | 130.5 | 131.4 | 383.0 | 128.9 |
| Pagrybis cemetary | 157 | 540–640 CE | 124.3 | 125.6 | 266.4 | 81.6 |
| Pagrybis cemetary | 207 | 565–654 CE | 121.8 | 123.3 | 249.5 | 80.6 |
| Taurapilis barrow cemetery | 1 | 3-6th c. CE | 126.9 | 128.0 | 302.6 | 88.3 |
| Taurapilis barrow cemetery | 4 | 259–538 CE | 129.5 | 130.4 | 310.3 | 90.5 |
| Marvelė cemetery | 113 | 134–408 CE | 131.8 | 132.6 | 347.4 | 94.4 |
| Taurapilis barrow cemetery | 5 | 236–530 CE | 124.1 | 125.5 | 269.8 | 82.2 |
| | *n* | | 8 | 8 | 8 | 8 |
| | Mean | | 127.4 | 128.5 | 306.6 | 92.1 |
| | Std. | | 3.7 | 3.4 | 44.8 | 15.7 |
| | Min. | | 121.8 | 123.3 | 249.5 | 80.6 |
| | Max. | | 131.8 | 132.6 | 383.0 | 128.9 |

**Table S17.** Estimated withers and croup heights, body masses and maximum rider weights (XRW) of Lithuanian horses dated to the Viking Age. Heights provided by regression equations in millimeters are converted to centimeters. Body mass and rider weight are in kilograms.

| Site | Specimen | Date | Withers height (*caballus*) | Croup height (*caballus*) | Body mass | XRW |
|---|---|---|---|---|---|---|
| Degsnė Labotiškė barrow cemetery | 5 | 9-11th c. CE | 121.6 | 123.1 | 268.4 | 84.1 |
| Marvelė cemetery | 35 | 9-11th.c. CE | 122.4 | 123.9 | 259.9 | 88.3 |
| Marvelė cemetery | 40 | 9-11th.c. CE | 134.6 | 135.1 | 374.0 | 114.3 |
| Marvelė cemetery | 49 | 9-11th.c. CE | 117.1 | 119.0 | 222.7 | 77.4 |
| Marvelė cemetery | 72 | 9-11th.c. CE | 126.7 | 127.8 | 282.9 | 87.7 |
| Marvelė cemetery | 103 | 9-11th.c. CE | 123.6 | 124.9 | 243.9 | 72.8 |
| Marvelė cemetery | 108 | 9-11th.c. CE | 130.0 | 130.9 | 294.3 | 80.6 |
| Marvelė cemetery | 116 | 9-11th.c. CE | 125.4 | 126.7 | 283.0 | 83.7 |
| Marvelė cemetery | 117 | 9-11th.c. CE | 119.3 | 121.0 | 257.7 | 83.4 |
| | *n* | | 9 | 9 | 9 | 9 |
| | Mean | | 124.5 | 125.8 | 276.3 | 85.8 |
| | Std. | | 5.4 | 5.0 | 42.6 | 11.7 |
| | Min. | | 117.1 | 119.0 | 222.7 | 72.8 |
| | Max. | | 134.6 | 135.1 | 374.0 | 114.3 |

**Table S18.** Estimated withers and croup heights, body masses, and maximum rider weights (XRW) of Lithuanian horses dated to the medieval period. Heights provided by regression equations in millimeters have been converted to centimeters. Body mass and rider weight are given in kilograms.

| Site | Specimen | Date | Withers height (*caballus*) | Croup height (*caballus*) | Body mass | XRW |
|---|---|---|---|---|---|---|
| Vilnius Lower castle | 67848 | 13-14th c. CE | 139.5 | 139.5 | 344.8 | 98.2 |
| Vilnius Lower castle | 14532 | 13-14th c. CE | 129.1 | 130.0 | 270.0 | 81.2 |
| Vilnius Lower castle | 66188 | 13-14th c. CE | 114.3 | 116.4 | 206.6 | 69.9 |
| Vilnius Lower castle | 66189 | 13-14th c. CE | 118.1 | 119.9 | 220.2 | 66.6 |
| Vilnius Lower castle | 4796 | 13-14th c. CE | 109.8 | 112.3 | 194.4 | 63.4 |
| Vilnius Lower castle | 66191 | 13-14th c. CE | 119.8 | 121.5 | 231.8 | 68.1 |
| Kernavė medieval town | K24 | 13-14th c. CE | 126.1 | 127.3 | 253.7 | 72.0 |
| Kernavė medieval town | K30 | 13-14th c. CE | 142.5 | 142.4 | 385.0 | 105.4 |
| Kernavė medieval town | K31 | 13-14th c. CE | 138.6 | 138.8 | 349.9 | 101.3 |
| Kernavė medieval town | K32 | 13-14th c. CE | 134.3 | 134.8 | 290.0 | 77.4 |
| Kernavė medieval town | K33 | 13-14th c. CE | 134.7 | 135.2 | 319.3 | 86.9 |
| Kernavė medieval town | K35 | 13-14th c. CE | 126.8 | 127.9 | 297.0 | 89.8 |
| Kernavė medieval town | K36 | 13-14th c. CE | 108.2 | 110.8 | 172.5 | 68.6 |
| Kernavė medieval town | K37 | 13-14th c. CE | 133.5 | 134.1 | 328.4 | 91.4 |
| Kernavė medieval town | K38 | 13-14th c. CE | 125.6 | 126.8 | 283.3 | 85.4 |
| Kernavė medieval town | K39 | 13-14th c. CE | 120.8 | 122.4 | 246.6 | 79.0 |
| Kernavė medieval town | K40 | 13-14th c. CE | 112.4 | 114.7 | 203.9 | 73.8 |
| Kernavė medieval town | K41 | 13-14th c. CE | 137.7 | 138.0 | 347.2 | 93.6 |
| Kernavė medieval town | K42 | 13-14th c. CE | 130.7 | 131.6 | 286.5 | 76.7 |
| Kernavė medieval town | K43 | 13-14th c. CE | 122.7 | 124.2 | 227.1 | 62.7 |
| Kernavė medieval town | K44 | 13-14th c. CE | 122.7 | 124.1 | 267.3 | 75.2 |
| Kernavė medieval town | K45 | 13-14th c. CE | 123.3 | 124.7 | 296.2 | 87.3 |
| Kernavė medieval town | K45.1 | 13-14th c. CE | 120.4 | 122.1 | 227.1 | 68.1 |
| Kernavė medieval town | K49 | 13-14th c. CE | 128.9 | 129.8 | 319.8 | 88.6 |
| Kernavė medieval town | K50 | 13-14th c. CE | 114.7 | 116.8 | 221.8 | 70.9 |
| Kernavė medieval town | K51 | 13-14th c. CE | 114.6 | 116.8 | 218.4 | 70.7 |

| | | | | | | |
|---|---|---|---|---|---|---|
| | n | | 26 | 26 | 26 | 26 |
| | Mean | | 125.0 | 126.3 | 269.6 | 79.7 |
| | Std. | | 9.6 | 8.8 | 56.1 | 12.1 |
| | Min. | | 108.2 | 110.8 | 172.5 | 62.7 |
| | Max. | | 142.5 | 142.4 | 385.0 | 105.4 |

**Table S19.** Withers heights (cm) of Lithuanian horses predating the Viking Age, estimated using equations introduced in this study, versus withers heights estimated following May (1985).

| **Site** | **Specimen** | **Date** | **Withers height (*caballus*)** | **Withers height (May 1985)** |
|---|---|---|---|---|
| Pagrybis cemetary | 142 | 6-7th c. CE | 130.3 | 130.1 |
| Pagrybis cemetary | 145 | 601–662 CE | 130.5 | 124.5 |
| Pagrybis cemetary | 157 | 540–640 CE | 124.3 | 127.3 |
| Pagrybis cemetary | 207 | 565–654 CE | 121.8 | 125.4 |
| Taurapilis barrow cemetery | 1 | 3-6th c. CE | 126.9 | 129.8 |
| Taurapilis barrow cemetery | 4 | 259–538 CE | 129.5 | 130.7 |
| Marvelė cemetery | 113 | 134–408 CE | 131.8 | 129.5 |
| Taurapilis barrow cemetery | 5 | 236–530 CE | 124.1 | 123.2 |
| | Mean | | 127.4 | 127.6 |
| | Std. | | 3.7 | 2.9 |
| | Min. | | 121.8 | 123.3 |
| | Max. | | 131.8 | 130.7 |

**Table S20.** Withers heights (cm) of Lithuanian horses dated to the Viking Age, estimated using equations introduced in this study, versus withers heights estimated following May (1985).

| Site | Specimen | Date | Withers height (*caballus*) | Withers height (May 1985) |
|---|---|---|---|---|
| Degsnė Labotiškė barrow cemetery | 5 | 9-11th c. CE | 121.6 | 119.0 |
| Marvelė cemetery | 35 | 9-11th.c. CE | 122.4 | 123.3 |
| Marvelė cemetery | 40 | 9-11th.c. CE | 134.6 | 133.9 |
| Marvelė cemetery | 49 | 9-11th.c. CE | 117.1 | 118.1 |
| Marvelė cemetery | 72 | 9-11th.c. CE | 126.7 | 126.9 |
| Marvelė cemetery | 103 | 9-11th.c. CE | 123.6 | 128.1 |
| Marvelė cemetery | 108 | 9-11th.c. CE | 130.0 | 132.2 |
| Marvelė cemetery | 116 | 9-11th.c. CE | 125.4 | 127.0 |
| Marvelė cemetery | 117 | 9-11th.c. CE | 119.3 | 120.4 |
| | n | | 9 | 9 |
| | Mean | | 124.5 | 125.4 |
| | Std. | | 5.4 | 5.6 |
| | Min. | | 117.1 | 118.1 |
| | Max. | | 134.6 | 133.9 |

**Table S21.** Withers heights (cm) of Lithuanian horses dated to the medieval period, estimated using equations introduced in this study, versus withers heights estimated following May (1985).

| Site | Specimen | Date | Withers height (*caballus*) | Withers height (May 1985) |
|---|---|---|---|---|
| Vilnius Lower castle | 67848 | 13-14th c. CE | 139.5 | 141.3 |
| Vilnius Lower castle | 14532 | 13-14th c. CE | 129.1 | 131.5 |
| Vilnius Lower castle | 66188 | 13-14th c. CE | 114.3 | 114.0 |
| Vilnius Lower castle | 66189 | 13-14th c. CE | 118.1 | 117.8 |
| Vilnius Lower castle | 4796 | 13-14th c. CE | 109.8 | 114.2 |
| Vilnius Lower castle | 66191 | 13-14th c. CE | 119.8 | 126.2 |
| Kernavė medieval town | K24 | 13-14th c. CE | 126.1 | 127.5 |
| Kernavė medieval town | K30 | 13-14th c. CE | 142.5 | 142.2 |
| Kernavė medieval town | K31 | 13-14th c. CE | 138.6 | 139.7 |
| Kernavė medieval town | K32 | 13-14th c. CE | 134.3 | 136.7 |
| Kernavė medieval town | K33 | 13-14th c. CE | 134.7 | 134.9 |
| Kernavė medieval town | K35 | 13-14th c. CE | 126.8 | 125.1 |
| Kernavė medieval town | K36 | 13-14th c. CE | 108.2 | 108.6 |
| Kernavė medieval town | K37 | 13-14th c. CE | 133.5 | 132.4 |
| Kernavė medieval town | K38 | 13-14th c. CE | 125.6 | 124.2 |
| Kernavė medieval town | K39 | 13-14th c. CE | 120.8 | 120.8 |
| Kernavė medieval town | K40 | 13-14th c. CE | 112.4 | 111.7 |
| Kernavė medieval town | K41 | 13-14th c. CE | 137.7 | 142.2 |
| Kernavė medieval town | K42 | 13-14th c. CE | 130.7 | 136.7 |
| Kernavė medieval town | K43 | 13-14th c. CE | 122.7 | 131.0 |
| Kernavė medieval town | K44 | 13-14th c. CE | 122.7 | 126.3 |
| Kernavė medieval town | K45 | 13-14th c. CE | 123.3 | 124.2 |
| Kernavė medieval town | K45.1 | 13-14th c. CE | 120.4 | 127.8 |
| Kernavė medieval town | K49 | 13-14th c. CE | 128.9 | 130.5 |
| Kernavė medieval town | K50 | 13-14th c. CE | 114.7 | 119.4 |
| Kernavė medieval town | K51 | 13-14th c. CE | 114.6 | 120.0 |
| | n | | 26 | 26 |
| | Mean | | 125.0 | 127.1 |
| | Std. | | 9.6 | 9.6 |
| | Min. | | 108.2 | 108.6 |
| | Max. | | 142.5 | 142.2 |

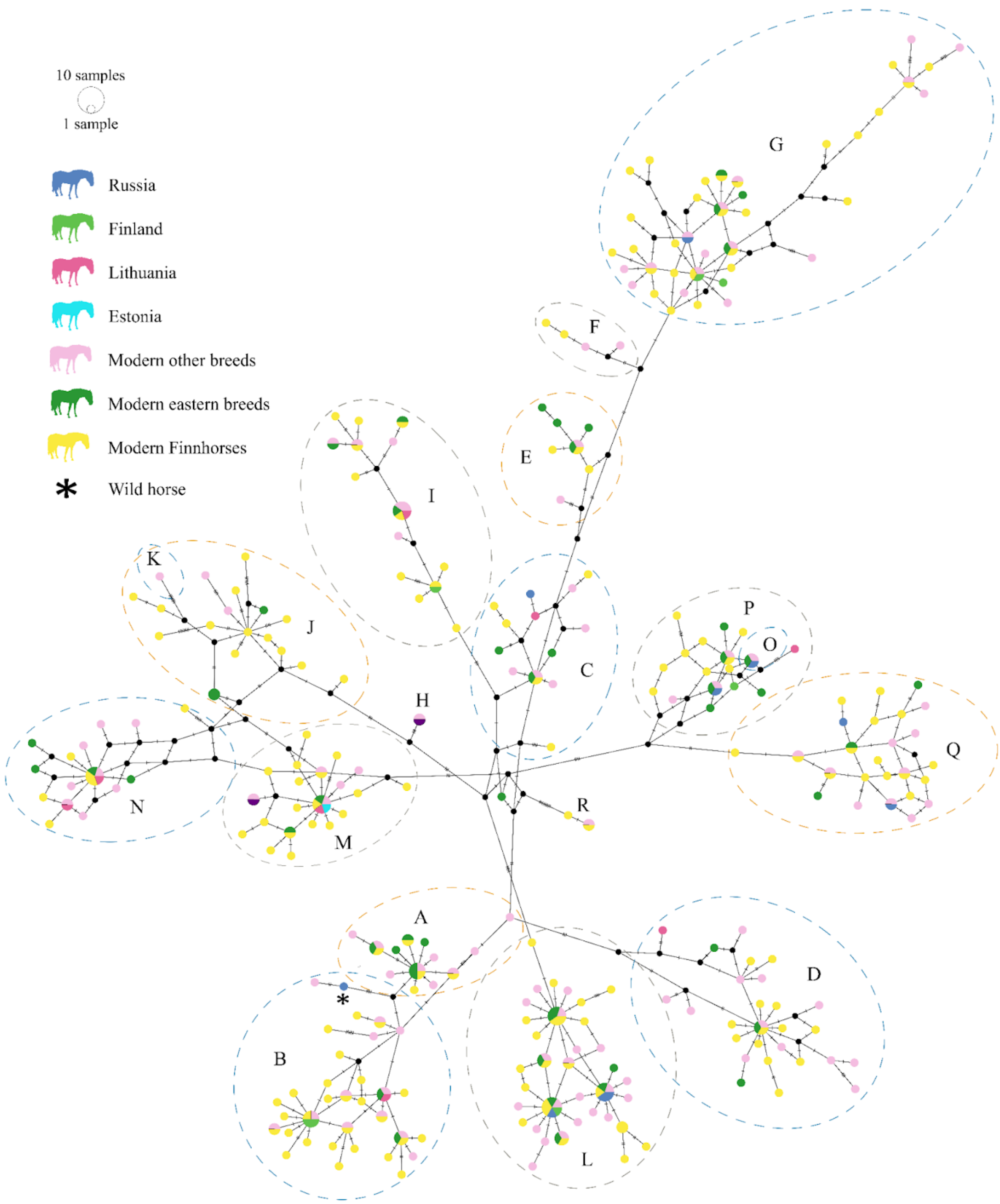


**Figure S1.** A median-joining haplotype network of 556 bp fragment of the mitochondrial control region of subfossil domestic horses and one wild horse (aEca44), along with modern horses used as reference samples. Modern horses were divided into three groups: Finnhorses, eastern breeds, and other horses, including sequences from Achilli et al. (2012) used to determine major haplogroups. Different countries of origin are presented with different colours. The size of each circle is proportional to the frequency of each haplotype, and tick marks across branches indicate mutational differences. The wild horse from Russia is marked with an asterisk.

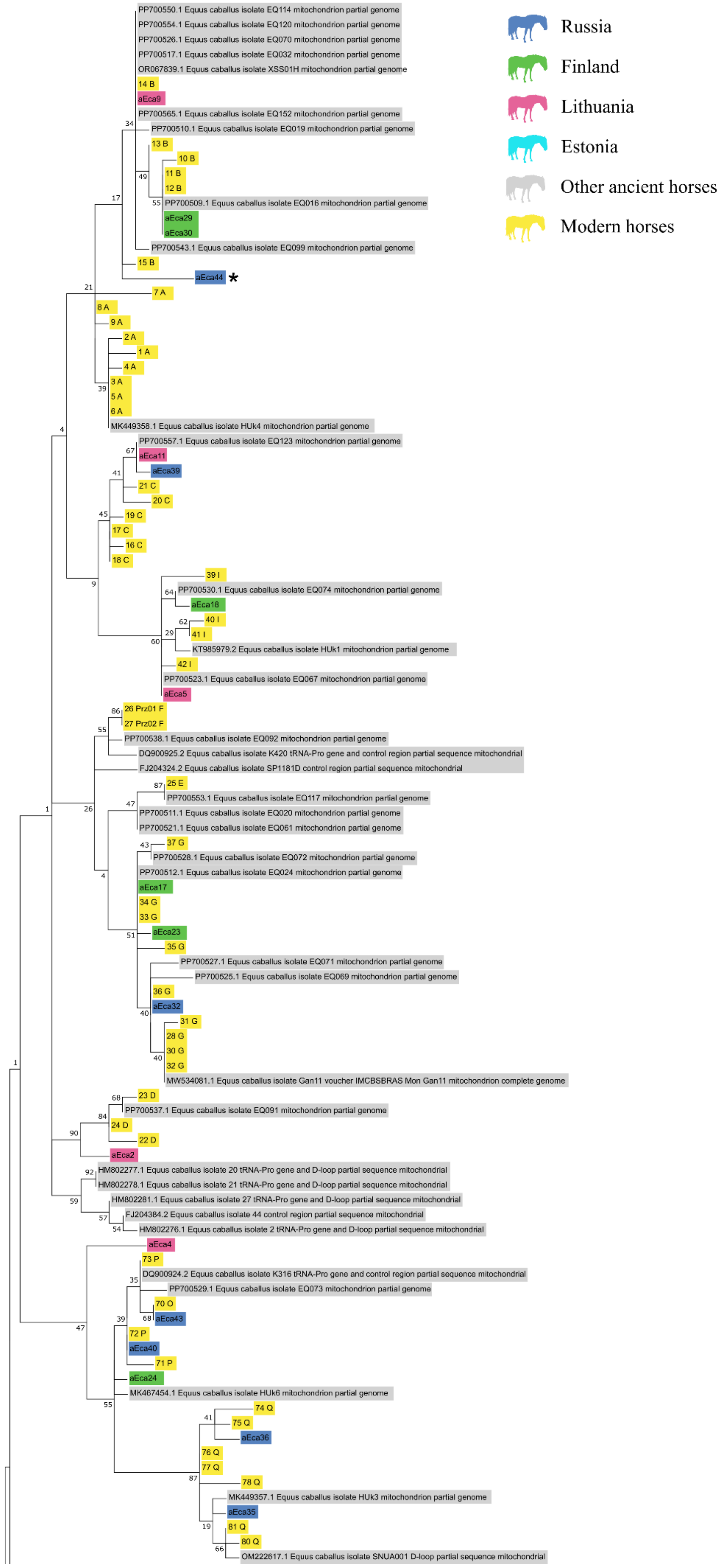

Russia
Finland
Lithuania
Estonia
Other ancient horses
Modern horses
PP700550.1 Equus caballus isolate EQ114 mitochondrion partial genome
PP700554.1 Equus caballus isolate EQ120 mitochondrion partial genome
PP700526.1 Equus caballus isolate EQ070 mitochondrion partial genome
PP700517.1 Equus caballus isolate EQ032 mitochondrion partial genome
OR067839.1 Equus caballus isolate XSS01H mitochondrion partial genome
14 B
aEca9
PP700565.1 Equus caballus isolate EQ152 mitochondrion partial genome
PP700510.1 Equus caballus isolate EQ019 mitochondrion partial genome
13 B
10 B
11 B
12 B
PP700509.1 Equus caballus isolate EQ016 mitochondrion partial genome
aEca29
aEca30
PP700543.1 Equus caballus isolate EQ099 mitochondrion partial genome
15 B
aEca44 *
7 A
8 A
9 A
2 A
1 A
4 A
3 A
5 A
6 A
MK449358.1 Equus caballus isolate HUk4 mitochondrion partial genome
PP700557.1 Equus caballus isolate EQ123 mitochondrion partial genome
aEca11
aEca39
21 C
20 C
19 C
17 C
16 C
18 C
39 I
PP700530.1 Equus caballus isolate EQ074 mitochondrion partial genome
aEca18
40 I
41 I
KT985979.2 Equus caballus isolate HUk1 mitochondrion partial genome
42 I
PP700523.1 Equus caballus isolate EQ067 mitochondrion partial genome
aEca5
26 Prz01 F
27 Prz02 F
PP700538.1 Equus caballus isolate EQ092 mitochondrion partial genome
DQ900925.2 Equus caballus isolate K420 tRNA-Pro gene and control region partial sequence mitochondrial
FJ204324.2 Equus caballus isolate SP1181D control region partial sequence mitochondrial
25 E
PP700553.1 Equus caballus isolate EQ117 mitochondrion partial genome
PP700511.1 Equus caballus isolate EQ020 mitochondrion partial genome
PP700521.1 Equus caballus isolate EQ061 mitochondrion partial genome
37 G
PP700528.1 Equus caballus isolate EQ072 mitochondrion partial genome
PP700512.1 Equus caballus isolate EQ024 mitochondrion partial genome
aEca17
34 G
33 G
aEca23
35 G
PP700527.1 Equus caballus isolate EQ071 mitochondrion partial genome
PP700525.1 Equus caballus isolate EQ069 mitochondrion partial genome
36 G
aEca32
31 G
28 G
30 G
32 G
MW534081.1 Equus caballus isolate Gan11 voucher IMCBSBRAS Mon Gan11 mitochondrion complete genome
23 D
PP700537.1 Equus caballus isolate EQ091 mitochondrion partial genome
24 D
22 D
aEca2
HM802277.1 Equus caballus isolate 20 tRNA-Pro gene and D-loop partial sequence mitochondrial
HM802278.1 Equus caballus isolate 21 tRNA-Pro gene and D-loop partial sequence mitochondrial
HM802281.1 Equus caballus isolate 27 tRNA-Pro gene and D-loop partial sequence mitochondrial
FJ204384.2 Equus caballus isolate 44 control region partial sequence mitochondrial
HM802276.1 Equus caballus isolate 2 tRNA-Pro gene and D-loop partial sequence mitochondrial
aEca4
73 P
DQ900924.2 Equus caballus isolate K316 tRNA-Pro gene and control region partial sequence mitochondrial
PP700529.1 Equus caballus isolate EQ073 mitochondrion partial genome
70 O
aEca43
72 P
aEca40
71 P
aEca24
MK467454.1 Equus caballus isolate HUk6 mitochondrion partial genome
74 Q
75 Q
aEca36
76 Q
77 Q
78 Q
MK449357.1 Equus caballus isolate HUk3 mitochondrion partial genome
aEca35
81 Q
80 Q
OM222617.1 Equus caballus isolate SNUA001 D-loop partial sequence mitochondrial

Figure continued from the next page.

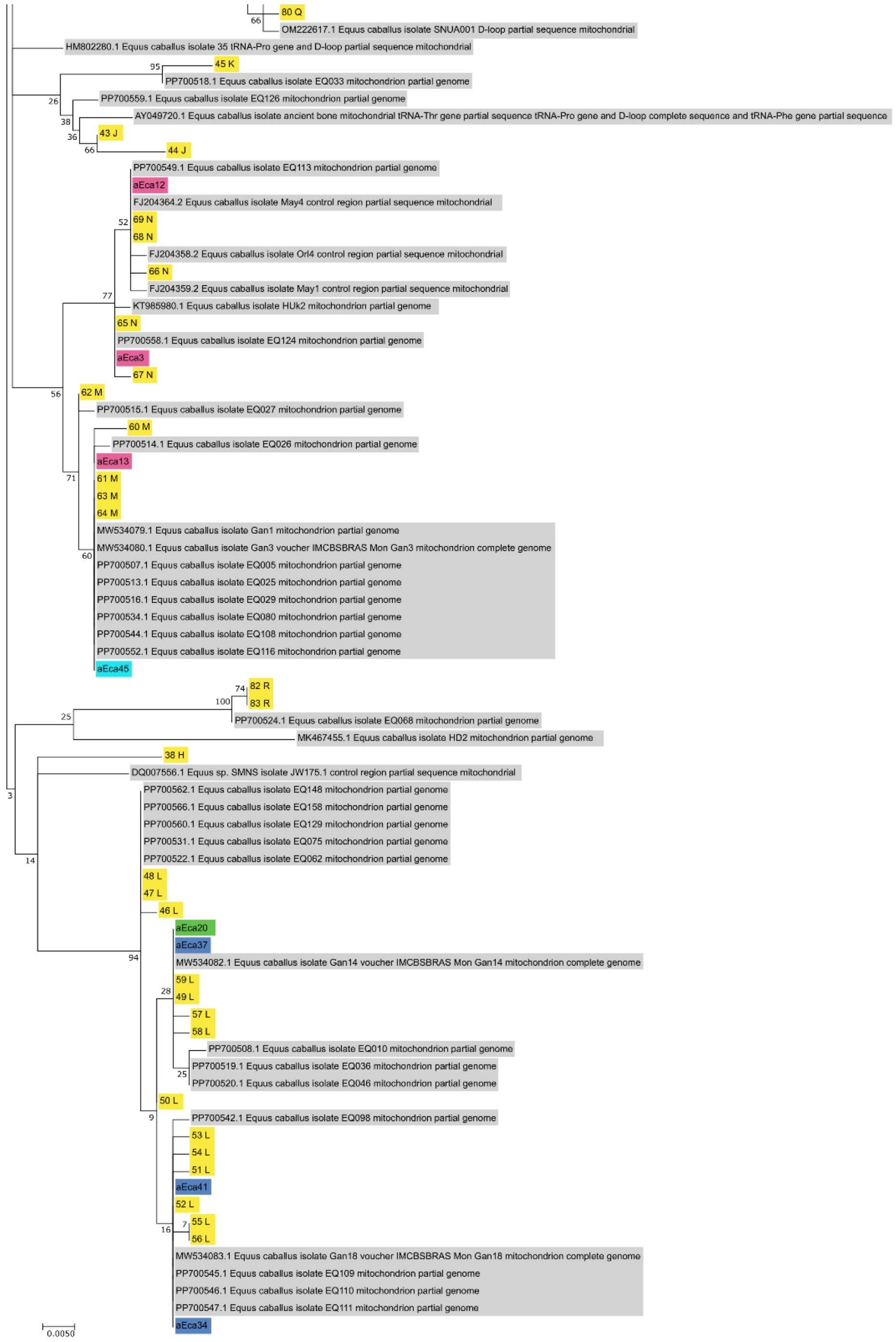


**Figure S2.** A maximum-likelihood phylogenetic tree of a 572-573 bp fragment of the mitochondrial control region of subfossil domestic horses and one wild horse (aEca44), ancient horses downloaded from GenBank, and 81 sequences from Achilli et al. (2012) used as a reference to determine the major haplogroups. Different countries of origin are presented with different colours. The wild horse from Russia is marked with an asterisk.
